# Reducing the Mitochondrial Oxidative Burden Alleviates Lipid-Induced Muscle Insulin Resistance in Humans

**DOI:** 10.1101/2023.04.10.535538

**Authors:** Matteo Fiorenza, Johan Onslev, Carlos Henríquez-Olguín, Kaspar W. Persson, Sofie A. Hesselager, Thomas E. Jensen, Jørgen F.P. Wojtaszewski, Morten Hostrup, Jens Bangsbo

**Affiliations:** August Krogh Section for Human Physiology, Department of Nutrition, Exercise and Sports, University of Copenhagen, Copenhagen, 2100, Denmark; Department of Biomedical Sciences, University of Copenhagen, Copenhagen, 2200, Denmark; August Krogh Section for Molecular Physiology, Department of Nutrition, Exercise and Sports, University of Copenhagen, Copenhagen, 2100, Denmark; Exercise Science Laboratory, Faculty of Medicine, Universidad Finis Terrae, Santiago 1509, Chile

**Keywords:** Insulin sensitivity, mitochondria, oxidative stress, reactive oxygen species (ROS), redox, mitochondrial function, bioenergetics, metabolism, diabetes

## Abstract

Pre-clinical models suggest a causative nexus between mitochondrial oxidative stress and insulin resistance. However, the translational and pathophysiological significance of this mechanism in humans remains unclear. Herein, we employed an invasive *in vivo* mechanistic approach in humans to manipulate mitochondrial redox state while assessing insulin action. To this end, we combined intravenous infusion of a lipid overload with intake of a mitochondria-targeted antioxidant (mtAO) in conjunction with insulin clamp studies. During lipid overload, insulin-stimulated muscle glucose uptake, as determined by the femoral arteriovenous balance technique, was increased by mtAO. At the muscle molecular level, mtAO did not affect canonical insulin signaling but augmented insulin-stimulated GLUT4 translocation while decreasing the mitochondrial oxidative burden under lipid oversupply. *Ex vivo* studies revealed that mtAO ameliorated features of mitochondrial bioenergetics, including diminished mitochondrial H_2_O_2_ emission, in muscle fibers exposed to high intracellular lipid levels. These findings provide translational and mechanistic evidence implicating mitochondrial oxidants in the development of lipid-induced muscle insulin resistance in humans.

**GRAPHICAL ABSTRACT:** 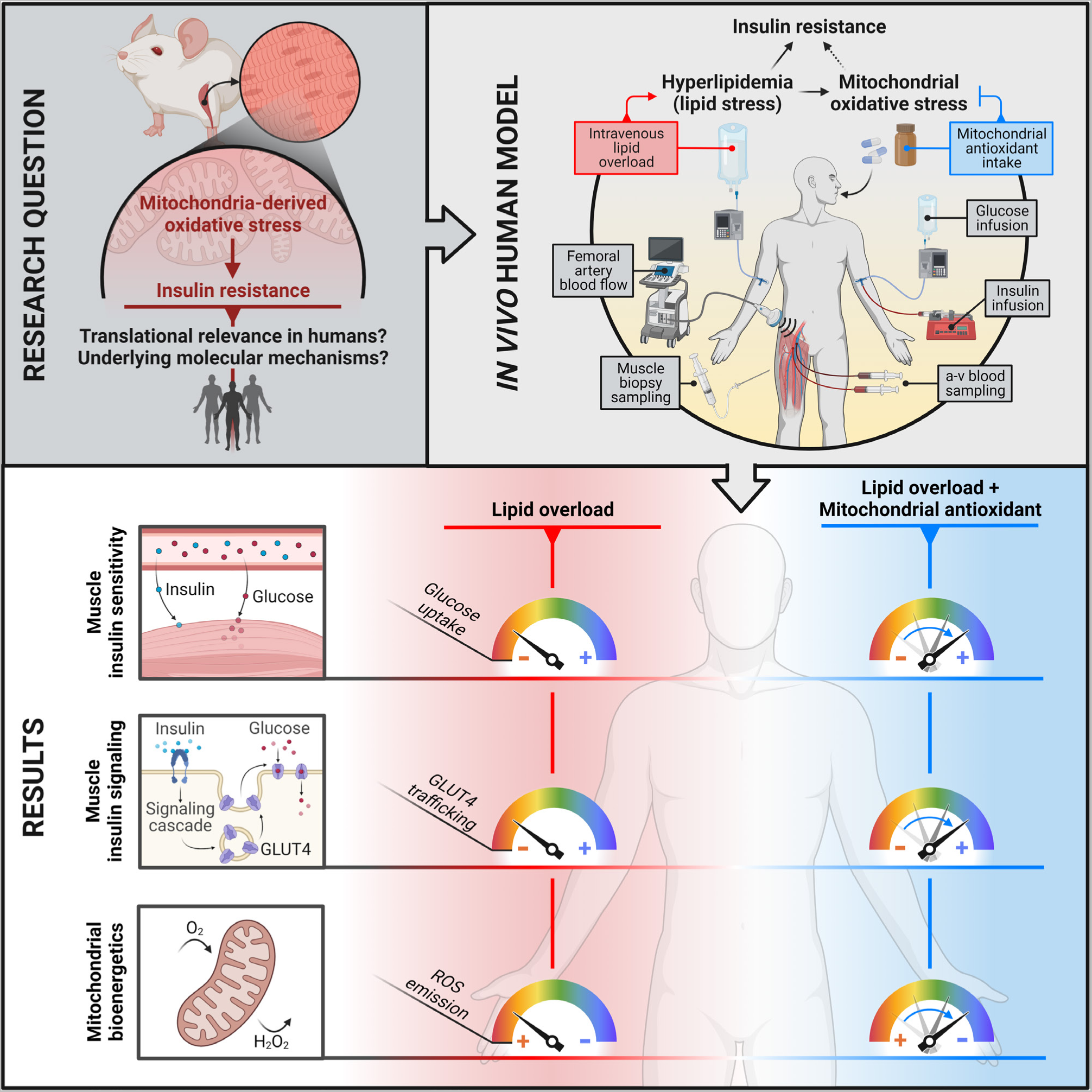

## INTRODUCTION

Insulin resistance, defined as defective responsiveness of peripheral tissues to the glucose-lowering action of insulin, is a main feature of type 2 diabetes. Skeletal muscle is the primary site of insulin-stimulated glucose uptake and thus the key tissue accounting for the impaired peripheral glucose disposal characterizing insulin-resistant states^1–3^.

Nutrient overload is a prominent cause of insulin resistance^4^. Particularly, excess fat intake may induce insulin resistance within skeletal muscle through the accumulation of intramyocellular lipids which ultimately suppress insulin signaling^5^. Acquired or inherited mitochondrial dysfunction has been proposed as a nexus between lipid oversupply and muscle insulin resistance^6^; a notion consistent with the impaired muscle mitochondrial respiratory function observed in individuals with obesity-related insulin resistance^7–9^. Mechanistically, dysfunctional mitochondria may display a reduced capacity to oxidize lipids, resulting in the accumulation of muscle lipid intermediates (i.e., acyl-CoAs, diacylglycerol/ceramides) that disrupt insulin signaling^10^. However, in contrast to such mitochondrial dysfunction theory of insulin resistance, genetic approaches that limit fatty acid uptake into muscle mitochondria, thereby inducing accumulation of cytosolic lipid intermediates, have been reported to improve whole-body glucose homeostasis^11–14^, suggesting that insulin resistance arises from excessive rather than diminished mitochondrial fat oxidation. Hence, although the mechanistic interplay between lipid oversupply, mitochondrial functions and insulin action remains controversial and incompletely understood^15–17^, pre-clinical data support an intertwined relationship between lipid-induced insulin resistance and mitochondrial stress^18^.

Mitochondria are a major source of superoxide and hydrogen peroxide (hereinafter referred to as reactive oxygen species (ROS))^19^. Excess ROS generation by mitochondria promotes a pro-oxidative shift in redox homeostasis leading to disrupted redox signaling and/or oxidative damage, i.e., oxidative stress^20^, which contributes to the pathogenesis of insulin resistance and diabetes^21, 22^. Because mitochondrial ROS production rates increase dramatically during fat oxidation^23–26^, this led to the supposition that mitochondria-derived oxidative stress is the connecting link between lipid oversupply and insulin resistance^18, 27^. Furthermore, independent of fat oxidation, mitochondrial oxidative stress may originate from the accumulation of cytosolic lipid intermediates, which, by inhibiting mitochondrial ADP transport, induce an increase in mitochondrial membrane potential and superoxide production^28, 29^. Collectively, the intracellular stresses imposed by lipid oversupply appear to converge upon mitochondria^30, 31^, with mitochondrial ROS likely acting as a critical signal in the cascade of events that impair insulin action. In support, genetic approaches to diminish mitochondrial ROS have been proven to effectively counteract insulin resistance induced by a high-fat diet in rodents^32–35^. This suggests that strategies targeting the mitochondrial oxidative burden holds therapeutic potential against insulin resistance under obesogenic conditions.

Mitochondria-targeted antioxidants are pharmacologic compounds that selectively deliver antioxidant moieties to mitochondria by utilizing either the mitochondrial membrane potential or affinity to a mitochondrial component^36–38^. Treatment with mitochondria-targeted antioxidant has been shown to preserve^32^ or ameliorate^39^ insulin action in high-fat diet-fed rodents and to convey greater metabolic benefits than inhibitors of non-mitochondrial ROS sources^39^.

Taken together, although pre-clinical data point towards a causative link between excess mitochondrial ROS and insulin resistance^40^, it is unknown whether this pathophysiological mechanism is conserved in humans. In this regard, clinical trials have reported inconsistent effects of general antioxidants on insulin action and glycemic control^41^; however, whether this is due to the unspecificity of general antioxidants to mitochondria remains elusive. Furthermore, species-specific differences in metabolism may contribute to translational barriers between pre-clinical models and humans^42^. Accordingly, human metabolic studies including selective targeting of mitochondrial redox state would be of high translational relevance and can help understand how mitochondrial ROS affect the molecular mechanisms controlling insulin action, ultimately aiding the development of new therapeutic strategies to counter insulin resistance.

We therefore conducted a proof-of-concept translational trial integrating invasive *in vivo* experiments in humans with *ex vivo* and *in vitro* studies in a variety of skeletal muscle models to investigate the mechanistic connection between muscle mitochondrial redox state and insulin sensitivity while interrogating the underlying mechanisms. Here, we show that mitochondrial oxidative burden reduction using a mitochondria-targeted antioxidant mitigates the impairments in muscle insulin sensitivity elicited by lipid oversupply, and that this occurs in association with rescue of the molecular machinery governing muscle insulin action along with improvements in mitochondrial bioenergetic function.

## RESULTS

### mtAO ameliorates lipid-induced insulin resistance in human skeletal muscle

To investigate the role of mitochondria-derived ROS in human skeletal muscle insulin resistance, we developed an integrative experimental model combining intravenous infusion of a lipid emulsion and heparin with intake of a mitochondria-targeted antioxidant (mtAO) to pharmacologically manipulate mitochondrial redox state *in vivo*. Through simultaneous hyperinsulinemic clamp studies, we assessed insulin-dependent muscle glucose uptake by means of the femoral arteriovenous balance technique (**Figure 1**A and B). Lipid plus heparin infusion was employed to acutely elevate plasma free fatty acid concentrations and progressively induce a lipotoxic environment in skeletal muscle, i.e. a metabolic state leading to insulin resistance^43, 44^ and increased mitochondrial ROS production^45^. To exacerbate the degree of insulin resistance and mitochondrial oxidative stress elicited by the lipid infusion, and thereby augment the potential salvage effect of mtAO, our mechanistic approach was implemented in overweight men with pre-diabetes (**Table S1**); a cohort that displays an insulin-resistant phenotype likely characterized by a pre-existing status of lipid-induced mitochondrial stress^46^.

**Figure 1.**
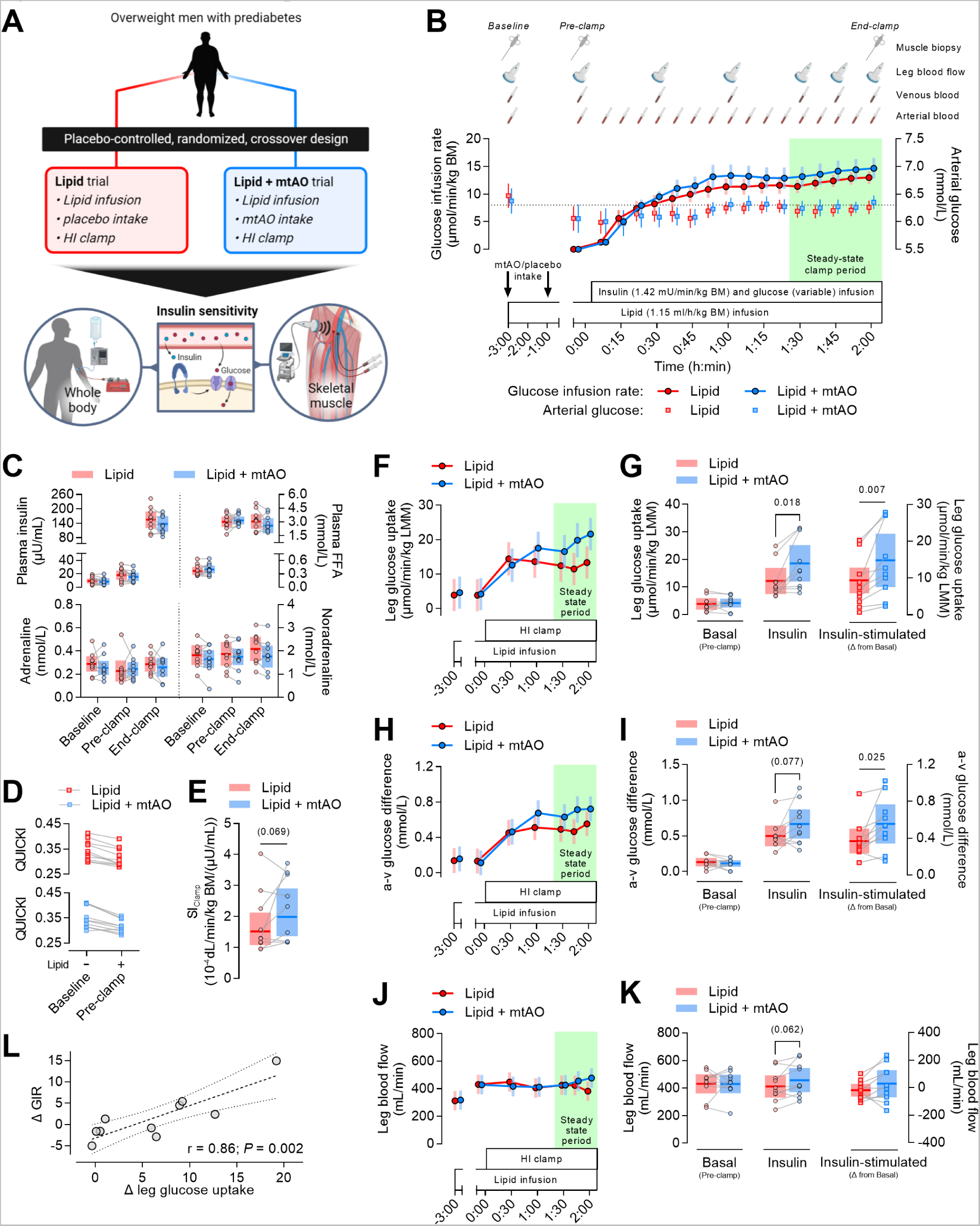
mtAO ameliorates lipid-induced insulin resistance in human skeletal muscle. (A) Schematic overview of the study design. HI clamp, hyperinsulinemic-isoglycemic clamp. (B) Schematic presentation of the experimental trial workflow, including the mean glucose infusion rate corrected per body mass (BM) and the arterial glucose concentrations. The dotted line indicates arterial glucose concentration corresponding to isoglycemia. Data are presented as means ± SEM. (C) Plasma insulin, free fatty acid (FFA) and catecholamine concentrations measured before the lipid infusion (Baseline), as well as before (Pre-clamp) and after (End-clamp) the HI clamp. Lipid + mtAO, n = 9 (End-clamp). (D) Individual changes in the quantitative insulin sensitivity check index (QUICKI) in response to 3 h of lipid infusion. (E) Whole-body insulin sensitivity expressed as the clamp-derived index of insulin sensitivity (SI_clamp_), defined as GIR/([Glu]×[ΔIns]), where GIR is the average glucose infusion rate during the steady-state clamp period, [Glu] is the average blood glucose concentration during the steady-state clamp period and [ΔIns] is the difference in plasma insulin concentration between baseline and the steady-state clamp period^131^. Lipid, n = 10; Lipid + mtAO, n = 9. (F-K) Skeletal muscle insulin sensitivity expressed as the leg glucose uptake, as calculated from the arteriovenous difference in plasma glucose concentration (determined by blood samples obtained from the femoral artery and the femoral vein) and the femoral arterial blood flow (determined by measurements of femoral artery internal diameter and blood flow velocity using ultrasound Doppler) corrected per leg muscle mass (LMM). (F, H and J) Time-course of leg glucose uptake, a-v glucose difference, and leg blood flow (data are estimated means ± 95% confidence limits). (G, I and K) Leg glucose uptake, a-v glucose difference, and leg blood flow before the HI clamp (Basal) and during the steady-state clamp period (Insulin; average from the last 30 min of the clamp). Between-treatment differences in insulin-stimulated leg glucose uptake, a-v glucose difference, and leg blood flow, calculated as the change (Δ) from the basal to the steady-state clamp period. (L) Pearson’s correlation between mtAO-induced changes in glucose infusion rate (GIR) and leg glucose uptake. Linear mixed models were used to estimate between-treatment differences (C, E, G, I and K). Data presented as observed individual values with estimated means ± 95% confidence limits, unless otherwise stated. n = 10, unless otherwise stated.

Lipid infusion markedly elevated circulating free fatty acids (**Figure 1C**) to an extent previously reported to impair insulin sensitivity at both the whole-body and skeletal muscle level^47–49^. The overall decrease in QUICKI observed after 3 h of lipid infusion confirmed the detrimental effect of the lipid overload on insulin sensitivity (**Figure 1D**).

To target the putative mitochondrial redox mechanisms underlying lipid-induced insulin resistance, capsules containing the mtAO mitoquinone mesylate or identical placebo capsules were orally administered before and during the lipid infusion in a randomized crossover fashion (**Figure S1** and **S2**C). Intake of mtAO ameliorated insulin action at the skeletal muscle level (**Figure 1**F and G), as measured by the insulin-stimulated glucose uptake across the leg, and non-significantly improved whole-body insulin sensitivity, as apparent from the 30% greater SI_clamp_ compared to placebo (**Figure 1**E). This was associated with non-significant trends toward greater arteriovenous glucose difference and femoral artery blood flow than placebo (**Figure 1**H-K). Thus, mtAO-dependent salvage of muscle glucose uptake was likely driven by increased peripheral glucose extraction in combination with enhanced glucose delivery through greater tissue perfusion; the latter aligning with previous observations of mtAO-induced improvements in vascular endothelial function^50, 51^, which is otherwise impaired by hyperlipidemia and may contribute to blunting the hyperemic response to insulin infusion^52, 53^.

Given the greater muscle glucose uptake elicited by mtAO under lipid overload, we sought to determine whether muscle substrate oxidation was differentially affected by mtAO as compared with placebo. We observed that, albeit a tendency toward a higher 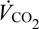 with mtAO (**Figure 2**A), neither muscle glucose nor lipid oxidation rates, as estimated by leg-specific indirect calorimetry^54^, were influenced by mtAO (**Figure 2**C and D). While recognizing the methodological limitations of the indirect calorimetry technique, our findings agree with the proposed dissociation between muscle insulin resistance and mitochondrial substrate oxidation^55^. To further interrogate the metabolic fate of glucose, we quantified lactate release across the leg, which has previously been shown to increase in response to lipid infusion^47, 48^. Leg lactate release was unaffected by mtAO (**Figure 2**E and F), reinforcing the absence of mtAO-dependent effects on substrate oxidation, and ultimately implying that mtAO improved muscle insulin sensitivity irrespective of increments in glucose oxidation.

**Figure 2.**
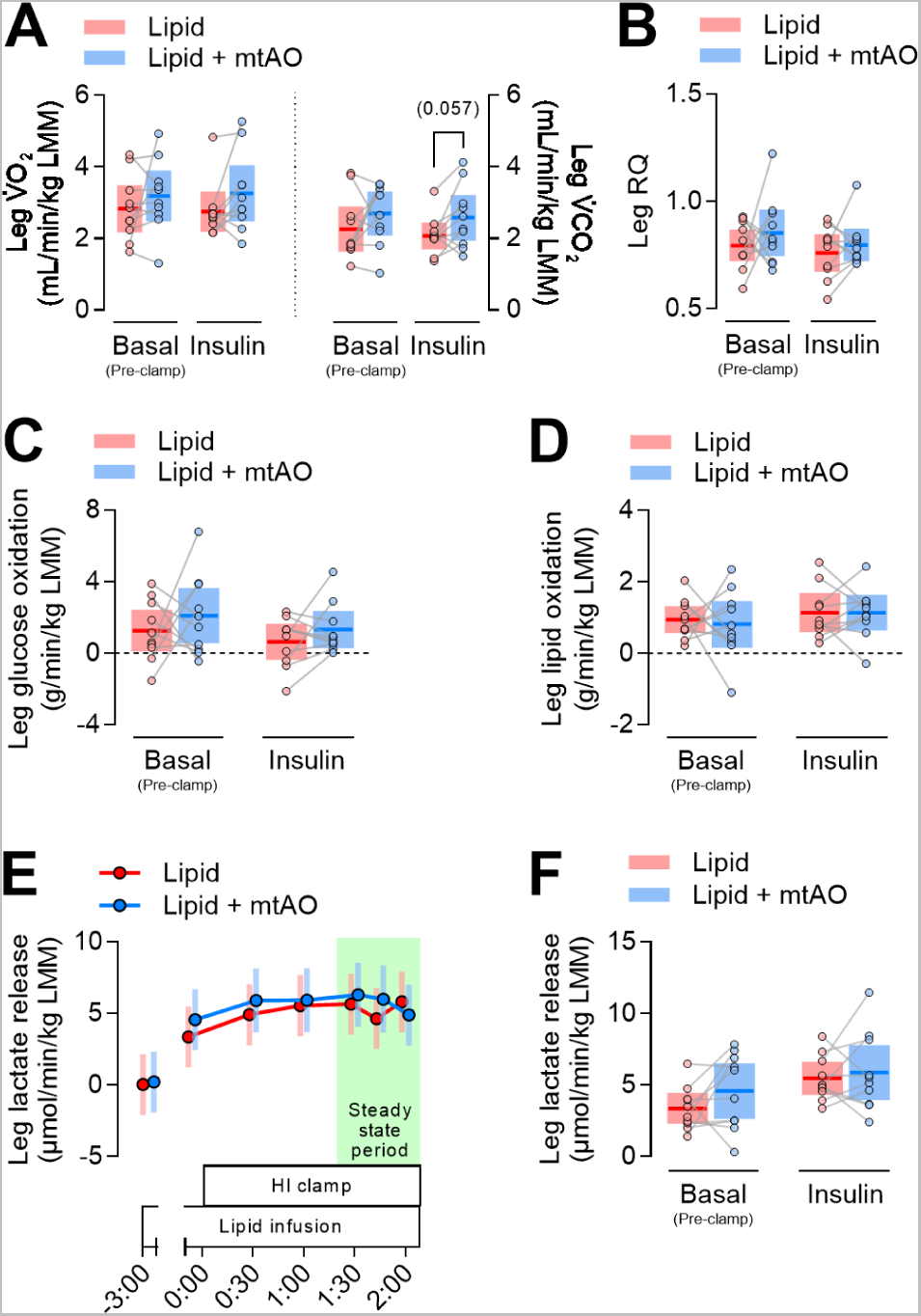
mtAO does not affect muscle substrate oxidation nor lactate release under lipid overload. (A-B) Leg O_2_ consumption 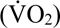, CO_2_ release 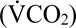 and respiratory quotient (RQ) before the HI clamp (Basal) and during the steady-state clamp period (Insulin). (C-D) Glucose and lipid oxidation across the leg, as estimated from the leg 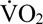 and 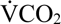, before the HI clamp (Basal) and during the steady-state clamp period (Insulin). (E-F) Leg lactate release as calculated from the arteriovenous difference in plasma lactate concentration and the femoral arterial blood flow. (E) Time-course of leg lactate release (data are estimated means ± 95% confidence limits). (F) Leg lactate release before the HI clamp (Basal) and during the steady-state clamp period (Insulin). Linear mixed models were used to estimate between-treatment differences (A, B, C, D and F). Data presented as observed individual values with estimated means ± 95% confidence limits, unless otherwise stated. n = 10.

Taken together, these data indicate that mtAO intake mitigated lipid-induced insulin resistance in human skeletal muscle *in vivo*, with this salvaging effect possibly driven by a synergistic increase in muscle glucose extraction and muscle tissue perfusion without apparent alterations in muscle substrate oxidation.

### mtAO does not affect canonical insulin signaling but enhances insulin-dependent GLUT4 translocation under lipid overload

Contrasting data have accumulated regarding the role of insulin signaling in lipid-induced insulin resistance. Some studies show impairments in proximal components of the insulin signaling pathway^56–59^ while others report intact insulin signaling^47, 48, 60–63^. To elucidate whether the observed insulin-sensitizing action of mtAO under lipid stress was associated with ameliorated insulin signaling, we assessed the phosphorylation of key proteins in the proximal and distal segments of the insulin signaling pathway (**Figure 3**A). We found that proximal insulin signaling at the level of Akt2 was unaffected by mtAO (**Figure 3**B), which is in contrast with the greater Akt phosphorylation observed during a glucose challenge in high-fat diet-fed rodents treated with the mtAO SS-31^32^. Likewise, distal insulin signaling, as measured by the phosphorylation of the Akt substrates GSK3 and TBC1D4, was unaffected by mtAO (**Figure 3**C and D).

**Figure 3.**
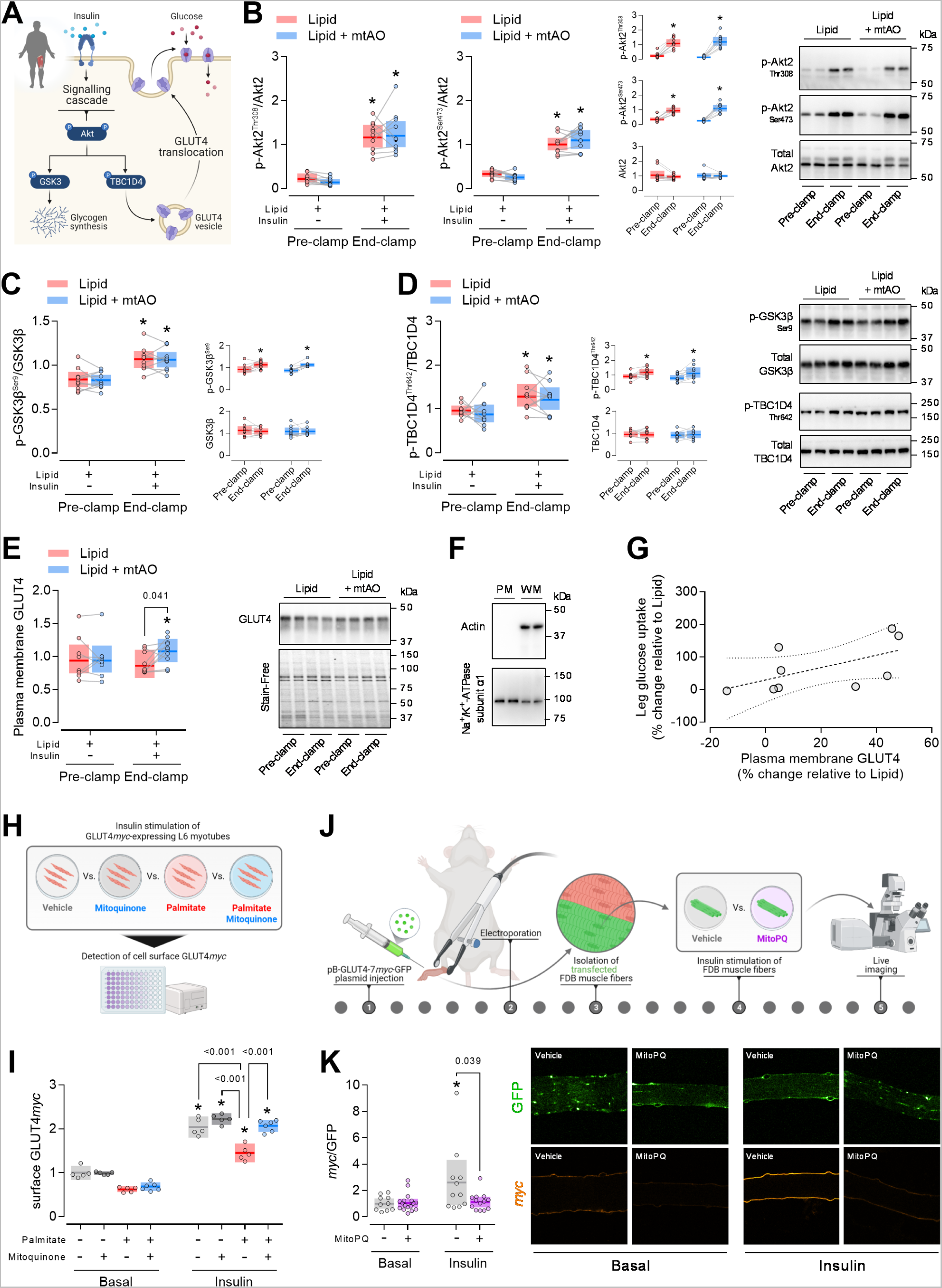
mtAO does not affect canonical insulin signaling but enhances insulin-dependent GLUT4 translocation under lipid overload. (A) Schematic overview of the major signaling events modulating insulin-stimulated glucose uptake in human skeletal muscle. (B) Proximal insulin signaling, as determined by phosphorylation of Akt2 on Thr308 and Ser473 in whole-muscle homogenates. (C-D) Distal insulin signaling, as determined by phosphorylation of the Akt substrates GSK3β on Ser9 (C) and TBC1D4 on Thr642 (D) in whole-muscle homogenates. (E) GLUT4 translocation, as determined by GLUT4 protein abundance in plasma membrane protein fractions. In-gel stain-free technology was used as a loading control. Lipid, n = 8 (Pre-clamp) and n = 9 (End-clamp); Lipid + mtAO, n = 10. Representative blots (n = 2 biological replicates from each muscle biopsy sample for each subject). (F) Purity of the isolated plasma membrane protein fraction, as determined by immunoblot analysis of actin (cytosolic protein marker) and Na^+^/K^+^-ATPase subunit α1 (plasma membrane protein marker) in plasma membrane homogenates (PM) as compared to the corresponding whole-muscle homogenates (WM). PM and WM samples were obtained by pooling a given volume of each individual sample analysed in B-F. (G) Pearson’s correlation between mtAO-induced changes in plasma membrane GLUT4 and leg glucose uptake under insulin stimulation. n = 9. (H) Workflow to determine insulin-stimulated GLUT4 translocation in L6 myotubes. (I) Insulin-stimulated GLUT4 translocation in L6 myotubes treated with either vehicle BSA (basal, n = 5; insulin, n = 5), or 50 nM mitoquinone (basal, n = 5; insulin, n = 5), or 250 µM palmitate (basal, n = 5; insulin, n = 5), or 250 µM palmitate + 50 nM mitoquinone (basal, n = 6; insulin, n = 6). Values are normalized to vehicle at basal. (J) Workflow to determine insulin-stimulated GLUT4 translocation in mouse FDB muscle fibers. (K) Insulin-stimulated GLUT4 translocation in mouse muscle fibers treated with either vehicle 0.1% ethanol (basal, n = 10; insulin, n = 11) or 10 µM MitoPQ (basal, n = 17; insulin, n = 13). Fibers were pooled from 3 mice. Values are normalized to vehicle at basal. Human data (B-E) are presented as observed individual values with estimated means ± 95% confidence limits. Linear mixed models were used to estimate within- and between-treatment differences at End-clamp. *Different from Pre-clamp (P<0.05). n = 10, unless otherwise stated. Rat muscle cell and mouse muscle fiber data (I and K) are presented as observed values with means ± 95% confidence limits. A one-way ANOVA was used to estimate between-treatment differences. *Different from “Basal” (P<0.05).

In light of pre-clinical data indicating that mitochondrial oxidative stress induces insulin resistance by impairing insulin-stimulated GLUT4 trafficking rather than canonical insulin signaling^34, 40^, we measured GLUT4 protein abundance in plasma membrane protein fractions as a proxy of GLUT4 translocation^64^. Insulin-mediated increments in sarcolemmal GLUT4 were apparent only with mtAO intake, which was associated with a greater sarcolemmal GLUT4 abundance than placebo (**Figure 3**E). As the changes in sarcolemmal GLUT4 were relatively small compared to the changes in muscle glucose uptake observed *in vivo* (**Figure 3**G), we conducted a series of experiments *in vitro* to further interrogate the role of mitochondrial ROS in muscle insulin resistance at the GLUT4 trafficking level. First, we assessed the effect of mtAO on GLUT4 translocation in L6 muscle cells expressing a *myc*-tagged GLUT4 (**Figure 3**H) and confirmed that mtAO rescued GLUT4 translocation defects elicited by palmitate treatment (**Figure 3**I). This suggests that mitochondrial ROS are necessary for lipid-induced impairments in GLUT4 trafficking. Next, using a genetically encoded biosensor GLUT4-7*myc*-GFP construct in mouse muscle^65^ (**Figure 3**J), we demonstrated that treatment with the mitochondria-targeted pro-oxidant Mito-Paraquat (MitoPQ) impaired insulin-mediated GLUT4 translocation (**Figure 3**K), revealing that mitochondrial ROS are sufficient to induce GLUT4-dependent insulin resistance in mature skeletal muscle.

Collectively, these findings indicate that mtAO did not affect canonical insulin signaling but enhanced insulin-stimulated GLUT4 trafficking in skeletal muscle under lipid stress.

### mtAO reduces the mitochondrial oxidative burden under lipid overload

To verify the link between the insulin-sensitizing and antioxidant action of mtAO under lipid stress, we measured markers of subcellular redox state in muscle biopsy samples obtained before initiation of the lipid infusion (Baseline) and upon termination of the HI clamp (End-clamp) (**Figure 4**A). Cytosolic and mitochondrial redox state were assessed by measuring peroxiredoxin 2 (PRDX2) and peroxiredoxin 3 (PRDX3) dimerization, respectively^66^. We found that, while the relative abundance of PRDX2 dimers was unchanged (**Figure 4**B), mtAO intake was associated with lower PRDX3 dimerization at the end of the HI clamp (**Figure 4**C), indicating a reduced mitochondrial oxidative burden compared to placebo. These findings support the compartmentalized action of mtAO and are consistent with pre-clinical data showing that the mitochondria-targeted pro-oxidant MitoPQ promotes dimerization of muscle PRDX3 without affecting PRDX2 redox state^40^.

**Figure 4.**
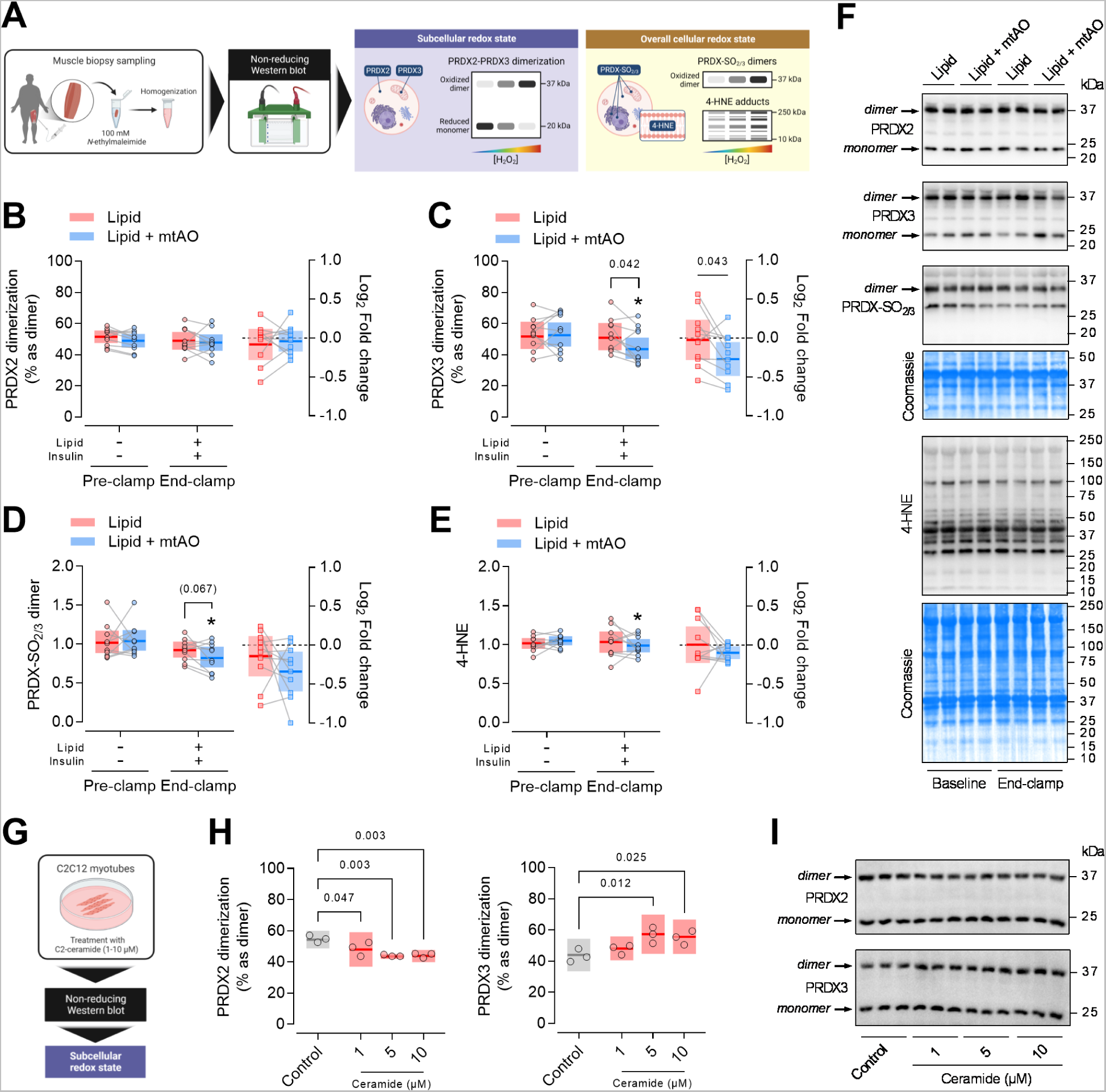
mtAO reduces the muscle mitochondrial oxidative burden under lipid overload. (A) Workflow to quantitatively analyse redox-sensitive proteins and lipid peroxidation in skeletal muscle biopsy samples. (B) Cytosolic oxidative burden as determined by protein abundance of peroxiredoxin 2 (PRDX2) dimers relative to monomers in whole-muscle homogenates. (C) Mitochondrial oxidative burden as determined by protein abundance of peroxiredoxin 3 (PRDX3) dimers relative to monomers in whole-muscle homogenates. (D) Overall peroxiredoxin oxidation as determined by protein abundance of oxidized/hyperoxidized peroxiredoxin (PRDX-SO_2/3_) dimers in whole-muscle homogenates. (E) Whole-muscle lipid peroxidation as determined by protein abundance of 4-hydroxynonenal (4-HNE) adducts in whole-muscle homogenates. (F) Representative blots related to the experiments in human skeletal muscle biopsy samples (n = 2 biological replicates from each muscle biopsy sample for each subject). Coomassie blue staining was used as a loading control for PRDX-SO_2/3_ and 4-HNE. (G-H) Cytosolic and mitochondrial oxidative burden in C2C12 myotubes treated with increasing levels of C2-ceramide. (I) Representative blots related to the experiments in C2C12 myotubes. Human data are presented as observed individual values with estimated means ± 95% confidence limits. Linear mixed models were used to estimate within and between-treatment differences. *Different from Baseline (P<0.05). n = 10 for all measurements. Mousce muscle cell data are presented as observed values with means ± 95% confidence limits. A one-way ANOVA was used to estimate between-treatment differences. n = 3 for all experiments.

To further characterize alterations in muscle redox state, we quantified the abundance of oxidized/hyperoxidized peroxiredoxin (PRDX-SO_2/3_) dimers, i.e. an indicator of overall peroxiredoxin oxidation^67, 68^, which tended to be lower with mtAO than with placebo (**Figure 4**D). Additionally, given that the mtAO mitoquinone exerts its main antioxidant action by preventing lipid peroxidation within the inner mitochondrial membrane^36, 69^, we determined the abundance of the lipid peroxidation product 4-hydroxynonenal (4-HNE) in whole-muscle homogenates. We found that, although mtAO intake was associated with a decrease in muscle 4-HNE from baseline, this did not correspond with lower 4-HNE levels compared to placebo (**Figure 4**E). This finding is consistent with data showing a lack of acute mitoquinone-dependent effects on blood lipid hydroperoxides^70^ and possibly relates to the selective action of mitoquinone on mitochondrial lipids, which constitute only a fraction of the myocellular lipidome and may, therefore, be overlooked when measuring lipid peroxidation at the whole-muscle level.

Interestingly, none of the readouts of muscle oxidative stress tended to be elevated in response to the 5-h lipid infusion (**Figure 4**B-E). This observation is in contrast with the increased skeletal muscle oxidative burden (as measured by the ratio of reduced to oxidized glutathione) reported upon an oral lipid overload^32^ and may be explained by the hyperinsulinemic state accompanying the lipid infusion in our study. Indeed, low-dose insulin has been reported to suppress ROS generation in human mononuclear cells^71^ and to exert an inhibitory action on oxidative stress in diabetic patients^72^. Furthermore, given that hyperinsulinemia *per se* may reduce systemic oxidative stress during hyperglycemic clamp conditions^73^, it is conceivable that the high-dose insulin infusion employed in the present study exerted an antioxidant action that abrogated lipid-induced increments in muscle oxidative stress.

To further interrogate the causal link between lipid overload and muscle mitochondrial oxidative stress, we conducted complementary studies in C2C12 muscle cells treated with increasing concentrations of ceramides (**Figure 4**G). Ceramide treatment induced a dose-dependent increase in PRDX3 dimers as opposed to a decrease in PRDX2 dimers (**Figure 4**H), thus corroborating the assumption that lipid stress promotes mitochondrial, but not cytosolic, oxidative stress in skeletal muscle.

Altogether, these data indicate that mtAO intake reduced the mitochondrial oxidative burden in human skeletal muscle under lipid overload, whose intrinsic pro-oxidant action was likely blunted by the concomitant hyperinsulinemic state.

### mtAO ameliorates muscle mitochondrial bioenergetics under lipotoxic conditions

To characterize lipid-induced alterations in human muscle mitochondrial function and interrogate the purported relieving action of mtAO, we conducted *ex vivo* analyses of muscle mitochondrial bioenergetics. To recapitulate the conditions characterizing the *in vivo* human experiments, mitochondrial bioenergetics was assessed in the presence of different intramyocellular lipid concentrations with and without pre-exposure to mtAO (**Figure 5**A and B). Specifically, either 20 µM or 60 µM palmitoyl-CoA (P-CoA), the long-chain fatty acid derived from palmitate, was employed to model a normal and a high intramyocellular lipid environment, respectively. Given the absence of *L*-carnitine, which is required for CPT1-dependent transport of P-CoA into the mitochondrial matrix, we studied lipid-induced mitochondrial stress driven by cytosolic lipid accumulation rather than accelerated fat oxidation. Indeed, cytosolic P-CoA, apart from promoting ROS generation via inhibition of mitochondrial ADP transport^28^, possesses amphiphilic properties that may alter the mitochondrial structure and cause peroxidation of the mitochondrial membrane^74, 75^; modifications which are supposedly prevented by the main antioxidant action of the mtAO mitoquinone.

**Figure 5.**
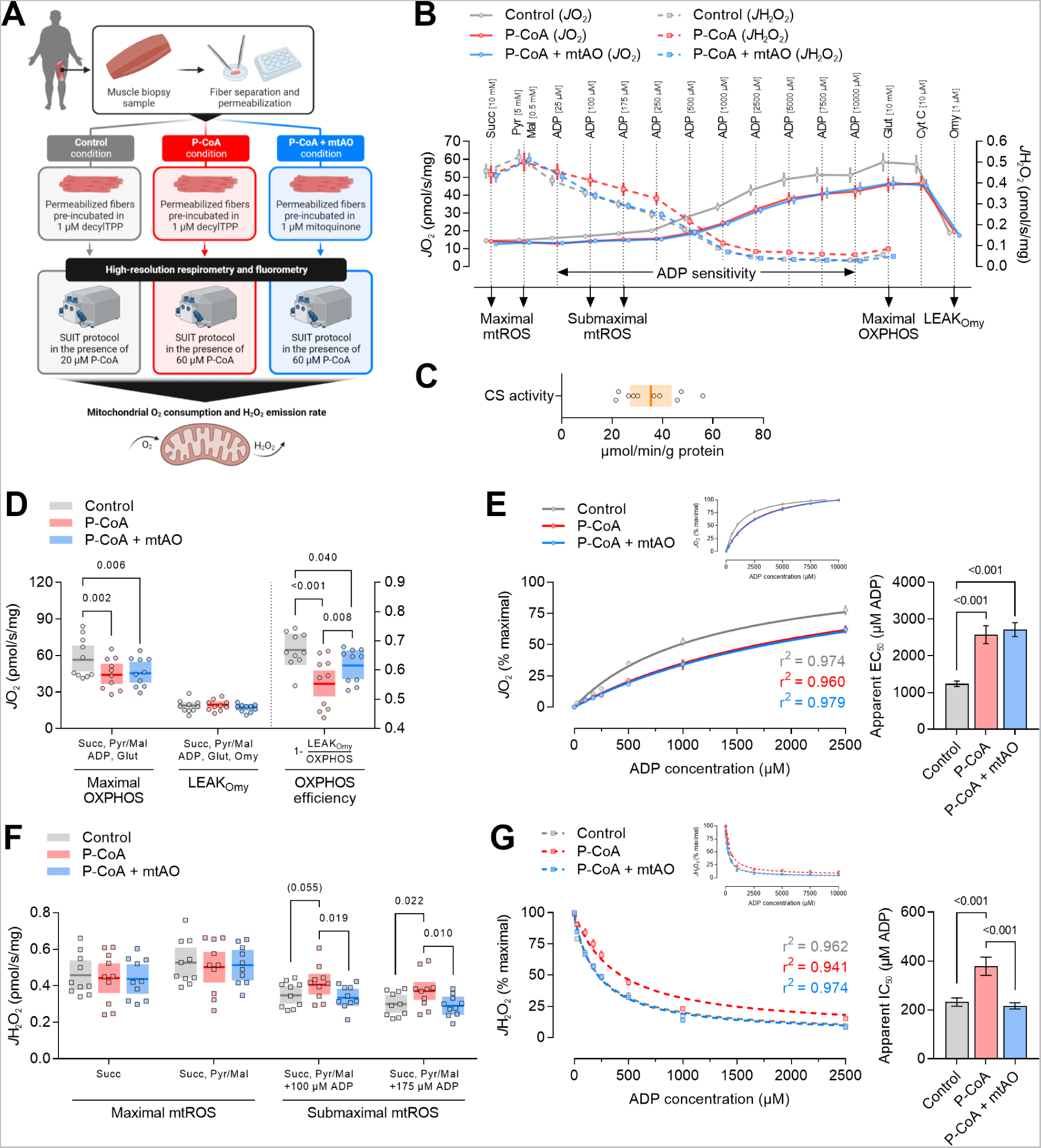
mtAO ameliorates muscle mitochondrial bioenergetics under lipotoxic conditions. (A) Workflow to determine *ex vivo* mitochondrial bioenergetics in skeletal muscle, i.e. in permeabilized muscle fibers, using a substrate-inhibitor titration (SUIT) protocol reflecting normal (20 µM P-CoA) vs. high (60 µM P-CoA) intramyocellular lipid conditions. Permeabilized muscle fibers were pre-incubated in either decylTPP (used as a control compound) or mitoquinone to evaluate the effects of mtAO on lipid-induced mitochondrial stress. (B) SUIT protocol employed to simultaneously measure mitochondrial O_2_ consumption (*J*O_2_) and H_2_O_2_ emission (*J*H_2_O_2_) rates. Succinate (Succ); pyruvate (Pyr); malate (Mal); adenosine diphosphate (ADP); glutamate (Glut); cytochrome C (Cyt C); oligomycin (Omy). Data are means ± SEM. (C) Muscle mitochondrial content, as determined by citrate synthase (CS) activity in muscle homogenates obtained from separate portions of the biopsy specimen used for assessments of mitochondrial bioenergetics. Data are presented as individual values with mean ± 95% confidence limits. (D) Maximal mitochondrial oxidative phosphorylation capacity (OXPHOS), oligomycin-induced leak respiration (LEAK_Omy_), and OXPHOS efficiency (calculated as 1-RCR = 1-LEAK_Omy_/OXPHOS^132^). Data are presented as individual values with mean ± 95% confidence limits. (E) Sensitivity of mitochondrial *J*O_2_ to ADP. The apparent half maximal effective concentration (EC_50_) for ADP was determined using [agoinst] vs. response (three parameters) analysis in GraphPad Prism. Data are means ± SEM. (F) Maximal and submaximal mitochondrial H_2_O_2_ emission rates. Data are presented as individual values with mean ± 95% confidence limits. (G) Sensitivity of mitochondrial *J*H_2_O_2_ to ADP. The apparent half maximal inhibitory concentration (IC_50_) for ADP was determined using [inhibitor] vs. response (three parameters) analysis in GraphPad Prism. Data are means ± SEM. A linear mixed model (D and F) or one-way ANOVA (E and G) was used to estimate between-treatment differences. n = 10 for all measurements.

We found that mitochondrial maximal oxidative phosphorylation capacity (OXPHOS) was impaired under lipid stress and was not restored by mtAO, whereas the leak respiratory state associated with OXPHOS (LEAK_Omy_) was unaffected by either lipid stress or mtAO (**Figure 5**D). In contrast, mtAO partly attenuated lipid stress-induced impairments in OXPHOS efficiency (**Figure 5**D); mechanistically, this may be due to mitoquinone-mediated changes in coenzyme Q redox status, which, by affecting mitochondrial supercomplex organization, can indirectly influence mitochondrial energy efficiency^76^. Notably, mitochondrial respiration appeared highly heterogeneous in the present cohort, possibly because of the large inter-individual variability in muscle mitochondrial content, as determined by citrate synthase activity (**Figure 5**C).

With respect to mitochondrial ROS, we observed that, under conditions that stimulate maximal mitochondrial ROS production (i.e., saturating concentrations of succinate in the absence of ADP), mitochondrial H_2_O_2_ emission rates were unaffected by either lipid stress or mtAO (**Figure 5**F), implying that ROS generated from complex I by reverse electron transport play a marginal role in lipid-induced mitochondrial stress. In contrast, under more physiological conditions stimulating submaximal ROS production (i.e., in the presence of ADP levels reflecting skeletal muscle at rest), we found that mitochondrial H_2_O_2_ emission rates were increased by lipid stress and that these increments were ablated by mtAO (**Figure 5**F). Accordingly, the effectiveness of mtAO in preventing excessive mitochondrial H_2_O_2_ emission in skeletal muscle was apparent only when the *in vitro* environment resembled the *in vivo* muscle milieu.

As mitochondrial OXPHOS dysfunction and excessive ROS production may arise from impaired sensitivity of mitochondria to ADP^29^, we also assessed whether lipid stress and/or mtAO altered such a feature of mitochondrial function. We found that the sensitivity of mitochondrial respiration (i.e., O_2_ consumption rate) to ADP was impaired under lipid stress and was not ameliorated by mtAO, matching prior clinical^28^ and pre-clinical data^29, 77^ (**Figure 5**E). On the other hand, mtAO restored the ability of ADP to suppress mitochondrial H_2_O_2_ emission, which was otherwise impaired under lipid stress (**Figure 5**G); a finding consistent with the beneficial effect of the mtAO SS-31 on the defective ADP sensitivity displayed by aging mouse muscle^78^.

Taken together, these data demonstrate that the impairments in muscle mitochondrial bioenergetics observed under lipid stress were partly rescued by mtAO. This was particularly evident in the rate of mitochondrial ROS emitted under experimental conditions resembling the *in vivo* metabolic environment of skeletal muscle at rest.

## DISCUSSION

Building on pre-clinical data indicating a causal role of excess mitochondrial ROS in the etiology of skeletal muscle insulin resistance, we designed an experimental model to interrogate the translational relevance of this mechanism in humans. By integrating pharmacological manipulation of mitochondrial redox state *in vivo* with simultaneous assessments of insulin-dependent glucose uptake into skeletal muscle, we demonstrated that intake of a mitochondria-targeted antioxidant (mtAO) enhanced muscle insulin sensitivity under lipid stress. This occurred in the absence of changes in canonical insulin signaling but in the presence of augmented insulin-mediated GLUT4 translocation along with reduced mitochondrial oxidative burden, as evidenced by complementary studies in human muscle biopsy samples, rodent muscle cells, and mouse muscle fibers. These results provide proof of principle that a pro-oxidative shift in mitochondrial redox state contributes to lipid-induced muscle insulin resistance in humans and suggest disrupted GLUT4 trafficking as one of the molecular mechanisms by which mitochondrial ROS affect insulin-stimulated glucose disposal.

In addition, through *ex vivo* experiments, we showed that a high intramyocellular lipid environment impaired features of mitochondrial bioenergetics in human muscle fibers and revealed that mtAO prevented lipid-induced increments in mitochondrial ROS emission. These data not only provide a comprehensive insight into the mitochondrial bioenergetic functions affected by lipid stress in human skeletal muscle but also demonstrate the effectiveness of mtAO in preserving some of these functions.

Our results contrast with the proposed irrelevance of mitochondrial ROS in the pathophysiology of muscle insulin resistance in humans^79^ and support findings from a seminal human intervention study inferring a role for mitochondrial oxidative stress in lipid-induced insulin resistance^32^. Specifically, Anderson and colleagues^32^ showed that muscle mitochondrial ROS production was acutely raised by a high-fat meal or a short-term high-fat diet; these human data, integrated with elegant rodent experiments, pointed toward a mechanistic link between lipid-induced mitochondrial ROS and insulin resistance in skeletal muscle. Nonetheless, the lack of complementary human experiments assessing muscle insulin action upon selective manipulation of mitochondrial redox state precluded drawing definitive conclusions on the translational significance of this mechanistic link in humans. Our model addresses this gap by incorporating insulin clamp-based measurements of muscle insulin action with intravenous infusion of a fat emulsion combined with intake of mtAO; a translational mechanistic approach which holds a number of methodological advantages. First, by applying the femoral arteriovenous balance technique to the gold standard insulin clamp technique, we were able to compute leg muscle-specific insulin sensitivity *in vivo*. Second, the use of intravenous instead of oral lipid overload enabled the induction of lipid stress without involving the gastrointestinal tract^80^, thus overcoming the insulinotropic effect of incretin hormones. Lastly, since lipid stress affects a multitude of cellular processes beyond mitochondrial redox homeostasis, the combination of mtAO intake with the intravenous lipid infusion permitted selective targeting of mitochondrial redox state.

The finding that lipid-induced insulin resistance was ameliorated by mtAO matches pre-clinical studies showing that genetic approaches to enhance mitochondrial ROS scavenging capacity can rescue insulin action under high-fat diet (HFD) feeding^32, 81–84^. In contrast, data from our human model partly contradict another pre-clinical study indicating that overexpression of mitochondria-targeted catalase (mCAT; a mitochondrial H_2_O_2_-specific scavenger) did not protect against insulin resistance in the acute lipid infusion model as opposed to the long-term HFD model of lipid overload^35^. This may be related not only to the markedly higher degree of lipid stress imposed by the acute lipid infusion in our human model than in the mCAT mouse model (i.e., 8-fold vs. 3-fold increase in circulating free fatty acids)^35^, but also to the differential mechanisms whereby mCAT and the mtAO mitoquinone exert their antioxidant action.

With respect to the metabolic benefits elicited by mitoquinone under lipid stress, our results align with those from studies in HFD-fed mice, where glycaemic control and glucose metabolism were improved by long-term treatment^39, 85–89^. On the other hand, evidence supporting the metabolic health-enhancing effects of mitoquinone in humans is scarce and limited to studies in metabolically healthy individuals, where long-term treatment did not lead to significant enhancements in glycaemic control indexes^51, 90^. However, it is noteworthy that the present results were obtained in an acute hyperlipidemic-hyperinsulinemic state that does not reflect normo-physiological conditions, implying that the favourable effects of mitoquinone on insulin action and glucose homeostasis may be apparent only in a setting of marked lipid-induced mitochondrial stress.

Skeletal muscle glucose uptake accounts for the vast majority of whole-body insulin-stimulated glucose disposal. Thus, under hyperinsulinemic clamp conditions, muscle insulin resistance is often accompanied by decreased rates of whole-body glucose disposal^5^. Our data indicate that the robust effect of mtAO on muscle insulin sensitivity was non-significant at the whole-body level, implying divergent effects in skeletal muscle and liver insulin sensitivity. Although an explanation for this is not apparent, a potential mechanism might involve redox-dependent regulation of Angiopoietin-like protein 4 (ANGPTL4), an endocrine factor that inhibits lipoprotein lipase^91^ and promotes the accumulation of hepatic tryglycerides in HFD-fed mice^92, 93^. Indeed, given that treatment with the antioxidant Tempol upregulates ANGPTL4 in mice under HFD feeding conditions^94^, it is tempting to speculate a similar effect of mtAO in the present lipid overload model, resulting in a greater fatty acid flux to the liver, which ultimately exacerbated the impairments in insulin-mediated suppression of hepatic glucose production.

Lipid oversupply may impair muscle insulin sensitivity via a multitude of mitochondria-dependent mechanisms, including i) mitochondrial substrate competition^95, 96^, ii) mitochondrial dysfunction-driven accumulation of intramuscular lipids^10^, and iii) excessive mitochondrial fatty acid oxidation^12^. In the present study, mtAO intake was not associated with alterations in either leg substrate oxidation or leg lactate release, suggesting a marginal role of mitochondrial substrate competition in the observed mitigation of lipid-induced muscle insulin resistance. Regarding intramuscular lipid accumulation, there is contrasting evidence for this phenomenon to occur upon acute lipid overload. Indeed, one study observed an increase in muscle ceramides^97^ as opposed to a number of studies showing no changes upon Intralipid infusion^47, 56, 59, 98^. Likewise, an Intralipid-induced increase in muscle diacylglycerol levels has been reported in some^56, 59^, but not all^47^, human studies. Intramuscular lipid accumulation has been proposed as a prominent cause of muscle insulin resistance in light of data showing that intramuscular lipids inhibit insulin signaling^56–58^. Since we did not observe mtAO-mediated alterations in insulin signaling, it is likely that, if intramuscular lipid accumulation occurred in the present study, mechanisms other than rescued insulin signaling contributed to alleviating insulin resistance. In this direction, myocellular accumulation of lipid intermediates may inhibit mitochondrial ADP transport, ultimately increasing mitochondrial membrane potential and ROS production^28, 29^; thus, it is plausible that mtAO attenuated the rise in mitochondrial oxidative burden associated with increased intramuscular lipid availability. Similarly, it is also likely that mtAO mitigated mitochondrial oxidative stress arising from the elevated β-oxidation associated with lipid oversupply.

Our finding that mtAO-dependent improvements in muscle insulin sensitivity occurred irrespective of enhancements in canonical insulin signaling, but in association with increased GLUT4 translocation, is in line with data showing that selective induction of mitochondrial ROS in muscle cells impairs insulin-stimulated GLUT4 translocation but not insulin signaling to Akt or Akt substrates^40^. Notably, the relatively small increase, or lack of change, in sarcolemmal GLUT4 observed upon insulin stimulation contrasts with the marked increase in muscle glucose uptake detected in either the presence or absence of mtAO; this was possibly due to the combination of two factors. First, the subcellular fractionation approach may underestimate GLUT4 translocation; indeed, in spite of a 100-fold increase in muscle glucose uptake, only a 2-fold increase in plasma membrane GLUT4 has been reported in response to exercise^99^. Second, the pre-diabetic state of our study participants might have further blunted the responsiveness of GLUT4 to insulin stimulation; in fact, increments in plasma membrane GLUT4 during a hyperinsulinemic clamp have been shown to be apparent in healthy but not in type 2 diabetic subjects^64^. Importantly, the present human results were substantiated by complementary experiments in rodent muscle cells, where mtAO rescued lipid-induced impairments in GLUT4 translocation, and rodent muscle fibers, where selective induction of mitochondrial oxidative stress impaired insulin-mediated GLUT4 translocation. Altogether, our findings indicate that a pro-oxidative shift in mitochondrial redox state is necessary and sufficient to disrupt GLUT4 translocation in skeletal muscle.

In the present study, the mtAO mitoquinone was employed as a tool to selectively target mitochondrial redox state. Mitoquinone is a ubiquinone moiety linked to a lipophilic cation, which enables it to pass through the plasma membrane and then accumulate into mitochondria driven by the plasma and the mitochondrial membrane potential, respectively^100^. Inside the mitochondrion, mitoquinone is absorbed to the mitochondrial inner membrane, where it acts as an antioxidant, primarily by preventing lipid peroxidation^101^. The mitoquinone-mediated protection of mitochondrial lipids from oxidative damage may indirectly reduce mitochondrial ROS production. Indeed, the mitochondrial inner membrane is rich in phospholipids very susceptible to ROS-induced peroxidation^102^, which may compromise the mitochondrial membrane structure and function, ultimately generating a chain reaction that induces a further increase in mitochondrial ROS^103^. In this scenario, mitoquinone may act as a chain-breaking antioxidant. Herein, through assessments of muscle subcellular redox state, we observed that mitoquinone reduced the mitochondrial oxidative burden, as measured by PRDX3 dimerization, without affecting the cytosolic redox state or the degree of lipid peroxidation when compared to placebo. While the lack of substantial effects on lipid peroxidation should be interpreted with caution (4-HNE adducts were quantified in whole muscle homogenates, possibly lacking the sensitivity to detect the mitochondria-specific alterations elicited by mitoquinone^104^), the present data demonstrate a mitochondria-specific antioxidant action of mitoquinone in human muscle under lipid stress; a finding that contrasts with a study reporting no effect of long-term mitoquinone treatment on age-related redox stress in rodent skeletal muscle^105^ and that possibly underlines the treatment duration-dependent effects of mitoquinone on muscle redox state.

The mitochondria-specific antioxidant effect of mitoquinone in human muscle was confirmed by assessments of mitochondrial bioenergetics *ex vivo*, where it prevented lipid-induced increments in mitochondrial H_2_O_2_ emission. These findings are consistent with the potential of mtAO to rescue impairments in muscle mitochondrial bioenergetics^78^. Notably, these effects were apparent only in the presence of substrates resembling the resting skeletal muscle metabolic milieu, thus pertaining to submaximal mitochondrial ROS production. Indeed, succinate-mediated maximal mitochondrial H_2_O_2_ emission rates were unaffected by either mtAO or lipid stress, which contrasts with pre-clinical data showing that treatment with the mtAO SS-31 reversed the increase in maximal mitochondrial H_2_O_2_ emission caused by a short-term high-fat diet^32^.

Overall, these outcomes not only underscore the importance of assessing mitochondrial bioenergetic function under biologically relevant conditions in human muscle^106–108^ but also imply that lipid-induced muscle mitochondrial stress is not driven by H_2_O_2_ originating from complex I via succinate-induced reverse electron transport (RET). This aligns with *in vitro* studies showing that fatty acids increase mitochondrial ROS generation under conditions of forward rather than reverse electron transport^109^. Similarly, our *ex vivo* muscle experiments did not show apparent effects of mitoquinone on RET-specific ROS production, as otherwise reported in rat heart mitochondria^110^. On the other hand, RET-specific ROS production was determined under saturating concentrations of succinate; thus, it cannot be ruled out that lipid stress- or mtAO-dependent alterations would have been apparent in the presence of non-saturating succinate levels.

Considering that, in muscle cells under resting conditions, RET (i.e., site I_Q_) and the outer quinone-binding site at complex III (site III_Q0_) are the main mitochondrial sources of total cellular H_2_O_2_ release, and together contribute to the vast majority of O_2_ ^-^/H O release in the mitochondrial matrix^111^, it is tempting to speculate that site III_Q0_ was the predominant mitochondrial source of ROS under lipid stress and that mitoquinone, by influencing the redox state of the coenzyme Q pool, indirectly attenuated lipid-induced H_2_O_2_ release from site III_Q0_. In this direction, the application of novel pharmacological tools, such as S1QEL and S3QEL molecules suppressing mitochondrial ROS production at specific sites of the electron transport chain^112, 113^, will allow to interrogate the role of given mitochondrial ROS sources in the pathophysiology of muscle insulin resistance.

## Conclusion and perspectives

This proof-of-concept study delineates a conserved role of mitochondria-derived ROS in the development of lipid-induced insulin resistance in human skeletal muscle. Altogether, our findings provide translational mechanistic insights into how mitochondrial biology affects muscle insulin action in humans and corroborate pre-clinical data proposing that mitochondrial oxidative stress causes insulin resistance by impairing insulin-stimulated GLUT4 translocation.

It is worth noting that muscle insulin resistance is multifactorial and may arise also from a lack of contractile activity. In this context, prolonged bed rest markedly reduces insulin sensitivity in the absence of concurrent alterations in muscle mitochondrial ROS production^114^, implying that alterations in mitochondrial redox state might be of secondary importance in inactivity-versus overnutrition-induced insulin resistance.

From a clinical standpoint, the present findings may underscore the therapeutic potential of mitochondrial redox targeting against obesity-related insulin resistance, thus providing a basis for larger scale trials addressing the clinical significance of mitochondria-targeted antioxidant therapy in the context of metabolic diseases.

## Limitations of the study

Owing to the invasiveness of the clinical procedures, the sample size of this proof-of-concept study was relatively small and restricted to a homogeneous cohort of adult men with pre-diabetes. The present subjects were chosen for being insulin-resistant but not yet treated with glucose- or lipid-lowering medications, which may affect muscle mitochondrial biology^115, 116^. Future studies should explore whether the mechanistic link between mitochondrial redox state and muscle insulin sensitivity is influenced by gender, age, and metabolic health status.

This study was designed to interrogate a putative mechanism underlying muscle-specific insulin resistance. Although comprehensive analyses were conducted at the skeletal muscle level, no assessments of hepatic insulin sensitivity or hepatic glucose metabolism were performed. Hence, it remains unclear whether selective targeting of mitochondrial redox state differentially affects muscle and liver insulin action. To interrogate the muscle tissue-specific delivery of orally administered mtAO, and thus infer its muscle-specific action, we conducted preliminary studies demonstrating that mitoquinone is taken up by skeletal muscle after administration of a single oral dose (**Figure S2**). However, given the cell type-specific effects of mitoquinone^117^, further studies are required to elucidate tissue-specific pharmacodynamic properties of mtAO in humans.

## ACKNOWLEDGMENTS

We thank all the study participants. We are grateful to Jens Jung Nielsen and Martin Thomassen (August Krogh Section for Human Physiology, Department of Nutrition, Exercise and Sports at the University of Copenhagen) for their technical support during the human experiments. We also thank Michael P. Murphy (MRC Mitochondrial Biology Unit, University of Cambridge) for the quantification of mitoquinone content in skeletal muscle biopsy samples.

This work was supported by a grant from the Novo Nordisk Foundation (grant number NNF18OC0052883). J.O. received funding from the Danish Diabetes Academy (grant number NNF17SA0031406). C.H.O. received funding from the Danish Diabetes Academy (grant number NNF17SA0031406).

## AUTHOR CONTRIBUTIONS

M.F. conceived the study, designed and conducted the experiments, analyzed data, and wrote the manuscript. J.O. contributed to designing and performing human experiments. C.H.O., K.W.P., S.A.H. and T.E.J. designed, carried out and analysed data from cell and animal experiments. J.F.P.W., M.H. and J.B. contributed to designing the human experiments and provided critical scientific advice. All authors contributed to data interpretation, critically revised the manuscript for important intellectual content and approved the final version of the manuscript.

## METHODS

### Key resources table

| REAGENT or RESOURCE | SOURCE | IDENTIFIER |
| --- | --- | --- |
| **Antibodies** |  |  |
| Phospho-Akt (Thr308) | Cell Signaling | Cat#9275 |
| Phospho-Akt (Ser473) | Cell Signaling | Cat#9271 |
| Akt2 | Cell Signaling | Cat#3063 |
| Phospho-GSK-3$\alpha/\beta$ (Ser21/9) | Cell Signaling | Cat#9331 |
| GSK-3β | BD Transduction Laboratories | Cat#610202 |
| Phospho-TBC1D4 (Thr642) | Cell Signaling | Cat#8881 |
| TBC1D4 | Abcam | Cat#ab189890 |
| GLUT4 | ThermoFisher | Cat#PA1-1065 |
| Na<sup>+</sup>/K<sup>+</sup>-ATPase subunit α1 | DSHB | Cat#α6F |
| Actin | Sigma-Aldrich | Cat#A2066 |
| Myc-Tag | Cell Signaling | Cat#2272 |
| PRDX2 | Abcam | Cat#ab109367 |
| PRDX3 | Abcam | Cat#ab73349 |
| PRDX-SO<sub>2/3</sub> | Abcam | Cat#ab16830 |
| 4-HNE | Alpha Diagnostics | Cat#HNE11-s |
| Goat Anti-Rabbit Ig, Human ads-HRP | SouthernBiotech | Cat#4010-05 |
| Goat Anti-Mouse Immunoglobulins/HRP | Agilent Technologies | Cat#P0447 |
| Alexa Fluor 568-donkey anti-rabbit IgG | Life Technologies | Cat#A10042 |
| **Chemicals, peptides, and recombinant proteins** |  |  |
| Lipid emulsion | Fresenius-Kabi | Intralipid® 20% 500 ml |
| Heparin (5000 U/ml) | LEO |  |
| Glucose solution (200 mg/ml) | Fresenius-Kabi |  |
| Human insulin | Novo Nordisk | Actrapid® 10 ml |
| DecylTPP | Santa Cruz Biotechnology | sc-264801 |
| Mitoquinone (human PmFBs experiments) | Michael P. Murphy, University of Cambridge | Gifted |
| Amplex UltraRed | Life Technologies | A36006 |
| Superoxide dismutase | Sigma | S 8160 |
| Blebbistatin | Sigma | B 0560 |
| Horseradish peroxidase | Sigma | P 8250 |
| Palmitoyl-CoA | Sigma | P 9716 |
| Succinate | Sigma | S 2378 |
| Malate | Sigma | M 1000 |
| ADP | Sigma | A 5285 |
| Glutamate | Sigma | G 1626 |
| Cytochrome C | Sigma | C 7752 |
| Oligomycin | Sigma | O 4876 |
| Antimycin A | Sigma | A 8674 |
| N-Ethylmaleimide | Sigma | E 1271 |
| Palmitate | Cayman Chemical | 29558 |
| Mitoquinone (L6 cell experiments) | MedChemExpress | HY-100116A |
| pB-GLUT4-7myc-GFP | Jonathan Bogan, Addgene | Plasmid #52872 |
| Mito-Paraquat | Abcam | ab146819 |
| C2-ceramide | Avanti Polar Lipids | 860502 |
| **Critical Commercial Assays** |  |  |
| Insulin ELISA kit | ALPCO | 80-INSHU-E01.1 |
| NEFA-HR(2) assay | Wako Chemicals |  |
| 2-CAT Plasma ELISA kit | LDN | BA E-4500 |
| Minute™ Plasma Membrane Protein Isolation Kit | Invent Biotechnologies Inc. | SM-005 |
| **Software and algorithms** |  |  |
| Datlab v.7.4 | Oroboros Instruments |  |
| GraphPad Prism v.9.5.0 | GraphPad |  |
| SAS Enterprise Guide 7.15 | SAS Institute Inc. |  |
| BioRender | BioRender |  |
| **Others** |  |  |
| Mitoquinone and placebo capsules | MitoQ Limited |  |

## RESOURCE AVAILABILITY

### Lead contact

Further information and requests for resources and reagents should be directed to and will be fulfilled by the lead contact, Matteo Fiorenza.

### Materials availability

This study did not generate new unique reagents.

### Data and code availability

Any additional information required to reanalyze the data reported in this paper is available from the lead contact upon request.

## EXPERIMENTAL MODEL AND SUBJECT DETAILS

### Human subjects

Twelve overweight men with prediabetes enrolled in the study. Participants were informed of risks and discomfort associated with the experimental procedures and provided written informed consent prior to inclusion in the study. One pariticipant withdrew from the study before the experimental trials due to major illness, whereas one participant was excluded during the first experimental trial due to anatomical incompatibility with the femoral catheters. Therefore, data from 10 participants who completed the study are reported here (Figure S1).

Inclusion criteria were: i) male, ii) age 40-60 years old, iii) body mass index 25-40 kg/m^2^, iv) fasting plasma glucose ≥5.6 mmol/L or HbA1c ≥5.7%, v) HOMA2-IR >1.4^118^, and vi) sedentary lifestyle (<3 h of physical activity/week and maximal oxygen uptake 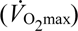 <45 mL/kg/min). Exclusion criteria were: i) current treatment with antidiabetic medications or insulin, ii) diagnosed thyroid, liver, heart or kidney disease iii) use of medications known to affect study outcome measures or increase the risk of study procedures, iv) use of mitoquinone or dietary supplements containing CoQ-10 within 30 days prior to the screening appointment, v) smoking or use of tobacco products, vi) recent weight gain or loss (>5% change within 2 months prior to inclusion in the study), and vii) excessive alcohol consumption (alanine aminotransferase >55 U/L). Subject characteristics are presented in **Table S1**.

The study was approved by the Regional Ethics Committee of Copenhagen, Denmark (H-19031551) and adheres to the principles of the Declaration of Helsinki. The study was registered at ClinicalTrials.gov (Identifier: NCT04558190).

### Animals

All animal experiments were approved by the Danish Animal Experimental Inspectorate (reference no. 2014-15-2934-01037) and complied with the European Union legislation outlined by the European Directive 2010/63/EU. To determine the effect of mitochondrial ROS on insulin-stimulated GLUT4 accumulation at the plasma membrane, the pB-GLUT4-7*myc*-GFP construct was expressed in skeletal muscle of male C57BL/6N mice at 12-14 weeks of age.

### Cell culture

L6 rat myoblasts stably expressing *myc*-tagged GLUT4 (L6-GLUT4*myc*) were grown to confluency in α-MEM (Gibco, 22571020) supplemented with 10% fetal bovine serum (FBS) (F0804; Sigma-Aldrich) and 1% Antibiotic-Antimycotic (Gibco, 15240062). The L6-GLUT4*myc* myoblasts generated by Prof. Amira Klip’s lab (SickKids Hospital, Toronto, Canada) and the C2C12 mouse myoblasts originally generated from Dr. Nobuharu L. Fujii’s lab (Tokyo Metropolitan University, Japan) were both kindly provided by Prof. Amira Klip. Muscle cells were maintained in a culture medium consisting of high-glucose Dulbecco’s Modified Eagle Medium (DMEM) (Cat. 11995073; Thermo Fisher Scientific) supplemented with 10% fetal bovine serum (F0804; Sigma-Aldrich) and 1% penicillin-streptomycin (Cat. 15140122; Thermo Fisher Scientific). L6 and C2C12 cells were kept at 37°C and in a humidified atmosphere of 5% CO_2_.

## METHOD DETAILS

### Screening visits

Potential participants underwent a pre-screening telephone interview followed by two in-person screening visits to assess eligibility criteria. The first screening visit comprised a complete medical history and a physical examination including lung and heart auscultation, electrocardiogram, blood pressure monitoring, and a fasting blood test. The second screening visit comprised assessments of body composition and cardiorespiratory fitness. Body composition parameters were determined using dual-energy X-ray absorptiometry (Lunar iDXA; GE Healthcare, GE Medical Systems, Belgium) after at least 4 h of fasting. Cardiorespiratory fitness 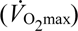 was determined during an incremental test to exhaustion on a mechanically braked cycle-ergometer (LC7; Monark Exercise AB, Vansbro, Sweden). The test protocol included a 4-min bout at 80 W followed by an incremental test with increments of 20 W/min until volitional exhaustion. Pulmonary gas exchanges were measured breath-by-breath using an on-line gas analysis system (Oxycon Pro, Viasys Healthcare, Hoechberg, Germany).

### Experimental trials

Participants attended two experimental trials on separate occasions interspersed by 14 days.

The experimental trials consisted of i) a hyperinsulinemic-isoglycemic clamp combined with intravenous infusion of a lipid emulsion and oral administration of placebo (“Lipid” trial), and ii) a hyperinsulinemic-isoglycemic clamp combined with intravenous infusion of a lipid emulsion and oral administration of mitochondria-targeted antioxidant (“Lipid + mtAO” trial).

Participants were instructed to maintain their usual lifestyle during the study period, i.e. individual interviews were conducted before each experimental trial day to ensure that no changes in dietary habits and physical activity routine occurred. Prior to the first experimental trial day, participants recorded food intake for three days, which was replicated prior to the second and third experimental trial days. Participants abstained from caffeine, alcohol, and exercise for 48 h prior to the experimental trial days.

### Administration of mitochondria-targeted antioxidant

Preliminary studies were conducted in a separate cohort of healthy young men (n = 6; age 24.5 ± 3.6 years, height 185 ± 10 cm, weight 87.5 ± 12.5 and 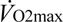 52.6 ± 4.7 mL/kg/min) to determine pharmacokinetic parameters of the mitochondria-targeted antioxidant mitoquinone in skeletal muscle after administration of a single oral dose (ClinicalTrials.gov Identifier: NCT04098510). Exclusion criteria were previous use of mitoquinone or CoQ-10 within 30 days of the screening appointment, ongoing treatment with medications, and smoking. Participants attended an experimental trial in the morning after an overnight fast and were instructed to abstain from caffeine, alcohol, and exercise for 24 h prior to the appointment. After 10 min of supine rest, a 3-mm incision was made over the lateral portion of the thigh under local anaesthesia and a biopsy (Pre dose) was obtained from the *vastus lateralis* muscle using a percutaneous Bergstrom needle with suction. Thereafter, participants ingested a single oral dose of mitoquinone [160 mg], corresponding to the maximal dose reported to be safe in humans^51^. Participants remained in the laboratory lying on a bed and were allowed to consume only water *ad libitum*. Additional biopsies were obtained from the *vastus lateralis* muscle at 1, 3, and 5 h after administration of mitoquinone (**Figure S2**A). Upon collection, muscle biopsy samples were washed in ice-cold saline to reduce blood contamination, dried, frozen in liquid N_2_, and stored in cryotubes at −80°C until analysis. Mitoquinone content in skeletal muscle was quantified by liquid chromatography-tandem mass spectrometry, as previously described^119^. None of the participants reported adverse events following mitoquinone administration.

The average maximum measured muscle concentration of mitoquinone (C_max_; 3.84 ± 1.10 ρmol/g) was observed 3 h after administration (**Figure S2**B). These data demonstrate that mitoquinone is taken up by human skeletal muscle *in vivo*, aligning with prior data showing mitoquinone levels of ∼20 ρmol/g in mouse muscle following long-term treatment with 500 μM mitoquinone^119^. It should be noted that the relatively low levels detected in humans as compared to mouse muscle are likely due to the prolonged treatment and the markedly higher dose administered in mice, i.e. ∼80 mg/kg, which corresponds to a human equivalent dose of 6.6 mg/kg^120^ as compared to the 1.6 mg/kg dose administered in our study. In light of this preliminary data, the main study included a short-term treatment period with escalating doses of mitoquinone to maximize muscle tissue accumulation and reduce any potential gastrointestinal discomfort associated with the high dose administered on the experimental trial day. Specifically, participants ingested daily doses of 40, 80, and 120 mg mitoquinone (or placebo) within the three days prior to the experimental trial day (**Figure S2**C). On the experimental trial day, a dose of 80 mg was administered 3 h and 1 h prior to the initiation of the hyperinsulinemic-isoglycemic clamp in order to achieve and maintain maximum muscle concentration of mitoquinone throughout the 120-min clamp period.

### Hyperinsulinemic-isoglycemic clamp

Participants reported to the laboratory in the morning after an overnight fast (∼10 h). Arrow catheters (20 gauge; Teleflex, Wayne, PA, USA) were inserted into the femoral artery and vein of the right leg under local anaesthesia (2 mL lidocaine without epinephrine, 20 mg/mL Xylocain) to obtain arterial and venous blood samples. Catheters were placed ∼3 cm below the inguinal ligament and advanced 10 cm in the proximal direction. Correct catheter placement was verified using ultrasound Doppler (Vivid E9; GE Healthcare, Waukesha, WI, USA). In addition, catheters were inserted into the antecubital veins for infusions. Thereafter, intravenous infusion of heparinized (0.2 units/min/kg body mass (BM)) lipid emulsion (1.15 mL/h/kg BM, Intralipid 20%) was initiated, and the first dose of mitoquinone [80 mg] or placebo was administered. After 2 h of lipid infusion, the second dose of mitoquinone [80 mg] or placebo was administered. After 3 h of lipid infusion, a hyperinsulinemic-isoglycemic clamp was initiated with a bolus of insulin (9.0 mU/kg BM) followed by 120 min of constant insulin infusion (1.42 mU/min/kg BM). During the clamp, glucose was infused from a 20% glucose solution. Arterial plasma glucose was measured every 7.5 min and the glucose infusion rate was adjusted to match isoglycemia, defined as the overnight-fasting arterial glucose concentration measured at baseline on the first experimental trial day. Femoral artery blood flow was measured, and blood was collected simultaneously from the femoral artery and femoral vein at −180, 0, 30, 60, 90, 105, and 120 min of the clamp (**Figure 1**B).

### Femoral artery blood flow

Femoral artery blood flow was determined by ultrasound Doppler (Vivid E9; GE Healthcare) equipped with a linear probe operating at an imaging frequency of 8.0 MHz and Doppler frequency of 3.1 MHz. Measurements were conducted in triplicate, i.e. three consecutive 15-s recordings were acquired at each time point, and the average value from the three replicate measurements was calculated.

### Blood and plasma analyses

Femoral arterial and venous blood samples were drawn in heparinized 2 mL syringes for immediate analysis of glucose, lactate, haematocrit, partial pressure of O_2_ (PO_2_) and CO_2_ (PCO_2_), O_2_ saturation, haemoglobin concentration, pH, and bicarbonate ions using an ABL800 Flex (Radiometer, Copenhagen, Denmark).

In addition, before the lipid infusion as well as before and after the hyperinsulinemic-isoglycemic clamp, an arterial blood sample was collected in 5 mL syringes and transferred to an Eppendorf tube containing 30 μL EDTA (0.2 M), after which it was centrifuged at 20,000 *g* for 3 min to collect plasma, which was stored at −80°C until analysis for plasma insulin, free fatty acids, and catecholamines. Plasma insulin concentration was measured using an enzyme-linked immunosorbent assay (ELISA) kit. Plasma free fatty acid concentration was measured using an enzymatic colorimetric assay adapted for the Pentra C400 (Horiba Medical). The kit reagents were reconstituted according to the manufacturer’s instructions, and the method was calibrated with a standard solution of oleic acid (1.0 mM) contained in the kit. Plasma adrenaline and noradrenaline concentrations were determined by using an ELISA kit.

### Leg muscle mass

The total muscle mass of the experimental leg was calculated as 83% of the leg lean mass, as measured by dual-energy X-ray absorptiometry (Lunar iDXA, GE Healthcare, GE Medical Systems, Belgium)^121^. The leg region, defined as the area extending from the inferior border of the ischial tuberosity to the distal tip of the toes, was adjusted using enCORE Forma v.15 software (GE Healthcare Lunar, Buckinghamshire, UK).

### Muscle biopsies

Muscle biopsies were obtained from the *vastus lateralis* muscle before lipid infusion (Baseline), as well as immediately before (Pre-clamp) and after (End-clamp) the hyperinsulinemic-isoglycemic clamp (**Figure 1**B). Biopsies were taken through separate incisions made over the lateral portion of the experimental thigh under local anaesthesia (1 mL lidocaine without epinephrine, 20 mg/mL Xylocain, AstraZeneca) using a percutaneous Bergström needle with suction.

Upon collection, muscle biopsy samples were washed with ice-cold saline to reduce blood contamination. The sampled biopsies were then divided in different portions, which were handled differently according to subsequent analyses. Specifically, a portion was snap-frozen in liquid N_2_ and stored at −80°C for subsequent immunoblotting and enzymatic activity analyses. Another portion was quickly immersed in PBS containing 100 mM *N*-ethylmaleimide (NEM; a fast-acting membrane permeable alkylating agent) for 5 min, snap-frozen in liquid N_2_, and stored at −80°C for subsequent non-reducing SDS-PAGE and Western blotting. A fraction (∼20 mg wet weight) of the biopsy specimen sampled at Baseline on the “Lipid” trial day was immediately placed in ice-cold biopsy preservation solution and prepared for assessments of mitochondrial bioenergetics.

### Immunoblotting

Protein abundance in whole-muscle homogenate lysates or plasma membrane fractions was determined by SDS-PAGE and Western blotting. Protein markers of muscle redox state (peroxiredoxin dimerization and 4-HNE) were resolved by SDS-PAGE under non-reducing conditions^122^.

#### Whole-muscle homogenate lysate preparation, SDS-PAGE and Western Blotting

Muscle samples were freeze-dried for 48 h and dissected free of blood, fat, and connective tissue. Dissection was performed under a stereo microscope with ambient temperature of ∼18°C and relative humidity <30%. After dissection, muscle tissue was weighed, and a fresh batch of ice-cold homogenization buffer (10% glycerol, 20 mM Na-pyrophosphate, 150 mM NaCl, 50 mM HEPES (pH 7.5), 1% Nonidet P-40 (NP-40), 20 mM β-glycerophosphate, 2 mM Na3VO4, 10 mM NaF, 2 mM PMSF, 1 mM EDTA (pH 8), 1 mM EGTA (pH 8), 10 µg/mL aprotinin, 10 µg/mL leupeptin, and 3 mM benzamidine) was added to the tube containing the freeze-dried tissue. 100 mM NEM was included in the homogenization buffer used for the samples undergoing non-reducing SDS-PAGE. Freeze-dried muscle samples (≈2 mg dry weight) were homogenized 2×2 min at 28.5 Hz in a TissueLyser (Qiagen TissueLyser II, Retsch GmbH, Haan, Germany Qiagen). Afterwards, the samples were rotated for 1 h at 4°C and sonicated (Branson Digital Sonifier) 2×10 s at 10% amplitude. The total protein concentration in each homogenate lysate sample was determined using a BSA standard kit (Thermo Scientific; assayed in triplicate). Each homogenate lysate sample was then mixed with 6×Laemmli buffer (7 mL of 0.5 M Tris base, 3 mL of glycerol, 0.93 g of DTT, 1 g of SDS, and 1.2 mg of bromophenol blue) and double-distilled H_2_O to reach equal protein concentration (2.0 µg/µL). DTT was not included in the buffer used for the samples subjected to non-reducing SDS-PAGE.

Equal amounts of total protein were loaded for each sample in precast gels (Criterion TGX Stain-Free Precast Gels, Bio-Rad, Copenhagen, Denmark). All the samples from each individual subject were loaded onto the same gel. Individual gels were compared using the same internal gel standard sample. Proteins were separated according to their molecular weight by SDS-PAGE and semi-dry transferred to a PVDF membrane (Millipore, Denmark). The membranes were blocked in either 2% skim milk or 3% BSA in Tris-buffered saline-Tween 20 (TBST) before being incubated overnight at 4°C in primary antibody diluted in either 2% skim milk or 3% BSA. After washing in TBST, the membranes were incubated with a secondary horseradish peroxidase-conjugated antibody for 1 h at room temperature. The secondary antibodies were diluted 1:5,000 in 2% skim milk or 3% BSA depending on the primary antibody. Membrane staining was visualized by incubation with a chemiluminescent horseradish peroxidase substrate (Immobilon Forte, Millipore, Denmark) before image digitalization on ChemiDoc MP (Bio-Rad). Western blot band intensity was determined by densitometry quantification (total band intensity adjusted for background intensity) using Image Lab Software v.6.0 (Bio-Rad Laboratories). For insulin signaling proteins, phosphorylated and total protein content were determined on separate membranes in separate analyses, and none of the membranes were stripped before protein quantification. Analyses were performed in duplicate for each biological sample; i.e. two separate samples were obtained from the same muscle specimen upon dissection, and the average signal intensity from the two samples was used as the result.

#### Plasma membrane fraction preparation, SDS-PAGE and Western Blotting

Plasma membrane proteins were isolated from frozen muscle samples (∼15 mg wet weight) using the Minute^TM^ Plasma Membrane Protein Isolation Kit (Invent Biotechnologies Inc.) according to the manufacturer’s instructions. The pellet containing plasma membrane proteins was dissolved in 30 µL of a fresh batch of ice-cold homogenization buffer (10% glycerol, 20 mM Na-pyrophosphate, 150 mM NaCl, 50 mM HEPES (pH 7.5), 1% Nonidet P-40 (NP-40), 20 mM β-glycerophosphate, 2 mM Na3VO4, 10 mM NaF, 2 mM PMSF, 1 mM EDTA (pH 8), 1 mM EGTA (pH 8), 10 µg/mL aprotinin, 10 µg/mL leupeptin, and 3 mM benzamidine). Each homogenate lysate sample was then mixed with 6×Laemmli buffer (7 mL of 0.5 M Tris base, 3 mL of glycerol, 0.93 g of DTT, 1 g of SDS, and 1.2 mg of bromophenol blue) and double-distilled H_2_O.

Equal volumes (10 µL) of plasma membrane homogenate lysates were loaded for each sample in precast gels (7.5% Criterion TGX Stain-Free Precast Gels, Bio-Rad). The samples from each individual subject were loaded onto the same gel. Individual gels were compared using the same internal gel standard sample. Proteins were separated according to their molecular weight by SDS-PAGE. Thereafter, the gels were activated by UV light exposure, and the total protein loaded for each sample was determined suing stain-free technology^123^. The activated gel was then electrophoretically transferred to a PVDF membrane. Membrane blocking and incubation were performed as described for whole-muscle homogenate lysates. Membrane staining was visualized by incubation with a high-sensitivity chemiluminescent horseradish peroxidase substrate (SuperSignal West Femto Maximum Sensitivity Substrate, Thermo Scientific) before image digitalization on ChemiDoc MP. Western blot band intensity was determined by densitometry quantification (total band intensity adjusted for background intensity) and normalized to the total protein loaded as determined by the stain-free method using Image Lab Software v.6.0 (Bio-Rad Laboratories). Analyses were done in technical duplicate, i.e. each sample was loaded twice in adjacent wells, and the average signal intensity was used as the result.

### Muscle mitochondrial bioenergetics

Mitochondrial O_2_ consumption and H_2_O_2_ emission (i.e., production minus removal) rates were simultaneously measured in permeabilized muscle fibers using high-resolution respirometry and fluorometry (Oxygraph O2k-FluoRespirometer; Oroboros Instruments, Innsbruck, Austria).

#### Preparation of permeabilized muscle fibers and pre-incubation with mtAO

Upon collection, muscle tissue was dissected free of connective tissue and fat. Multiple small muscle bundles (1.5-2.5 mg wet weight) were prepared from each muscle tissue sample. Each bundle was gently teased along the longitudinal axis with needle tip forceps in ice-cold biopsy preservation solution (2.77 mM CaK_2_-EGTA, 7.23 mM K_2_EGTA, 5.77 mM Na_2_ATP, 6.56 mM MgCl_2_, 20 mM taurine, 15 mM Na_2_Phosphocreatine, 20 mM imidazole, 0.5 mM DTT, 50 mM K-MES, pH 7.1). Teased fiber bundles were treated with 50 µg/mL saponin for 20 min under gentle rocking on ice to permeabilize the cell membranes while preserving the mitochondrial membranes. Permeabilized fiber bundles (PmFBs) were then transferred to mitochondrial respiration medium (buffer Z: 1 mM EGTA, 5 mM MgCl_2_, 105 mM K-MES, 30 mM KCl, 10 mM KH_2_PO_4_, 5 µM pyruvate, 2 µM malate, and 5 mg/mL BSA, pH 7.4) supplemented with either 1 µM decylTPP (i.e., the mitochondria-targeting moiety lacking the antioxidant group) or 1 µM mitoquinone, where they remained for 20 min under gentle rocking on ice; an incubation step allowing for accumulation of decylTPP/mitoquinone into mitochondria, driven by the mitochondrial membrane potential^124^. DecylTPP was used as a control compound to account for the nonspecific effects of mitoquinone on mitochondria^69^. Thereafter, PmFBs were transferred to a new batch of buffer Z without decylTPP/mitoquinone and rinsed by rocking for 10 min on ice. Then, PmFBs were gently blotted on dry filter paper for 15 s and weighed before being added to an Oxygraph O2k-FluoRespirometer chamber containing 2 mL of buffer Z supplemented with 25 µM blebbistatin, 10µM Amplex UltraRed, 5 U/mL superoxide dismutase, and 1 U/mL horseradish peroxidase.

#### Substrate-inhibitor titration protocol

Mitochondrial bioenergetics was determined using a substrate-inhibitor titration protocol validated for simultaneous measurements of mitochondrial O_2_ (*J*O_2_) and H_2_O_2_ (*J*H_2_O_2_) fluxes and ADP sensitivity in human PmFBs^108^. Experiments were conducted in the presence of palmitoyl-CoA (P-CoA) concentrations reflecting normal (20 µM) or high (60 µM) intracellular lipid conditions^29, 125^. Specifically, 20 µM P-CoA was titrated in the chambers containing PmFBs pre-treated with decylTPP (“Control” sample), whereas 60 µM P-CoA was titrated in the chambers containing PmFBs pre-treated with decylTPP (“P-CoA” sample) or mitoquinone (“P-CoA + mtAO” sample) (**Figure 5**A). The substrate-inhibitor titration protocol started with the addition of 10 mM succinate to induce reverse electron flow, followed by titration of 5 mM pyruvate and 0.5 mM malate to stimulate maximal mitochondrial H_2_O_2_ emission (Figure 4B). Thereafter, increasing amounts of ADP (25-10000 µM) were titrated stepwise to determine submaximal mitochondrial H_2_O_2_ emission rates and the sensitivity of mitochondrial O_2_ consumption and H_2_O_2_ emission to ADP. 10 mM glutamate was then titrated to determine maximal oxidative phosphorylation (OXPHOS) capacity through complex I and II, followed by titration of 10 µM cytochrome C to test mitochondrial outer membrane integrity. 1 µM oligomycin was then added to inhibit ATP synthase and measure oligomycin-induced leak respiration (LEAK_Omy_). Finally, 2.5 µM antimycin A was titrated to terminate respiration and allow for correction for non-mitochondrial oxygen consumption.

The sensitivity of mitochondrial O_2_ consumption (apparent EC_50_) and H_2_O_2_ emission (apparent IC_50_) to ADP was estimated using [agoinst] vs. response (three parameters) analysis and [inhibitor] vs. response (three parameters) analysis, respectively, in GraphPad Prism. The first concentration of ADP (25 µM ADP) was not used to estimate the apparent EC_50_ in “P-CoA” and “P-CoA + mtAO”, as this concentration of ADP did not stimulate respiration in the presence of 60 µM P-CoA.

Experiments were performed at a chamber temperature of 37°C. Instrumental and chemical O_2_ background fluxes were calibrated as a function of O_2_ concentration and subtracted from the total volume-specific O_2_ flux (Datlab v.7.4 software; Oroboros Instruments). O_2_ levels were maintained between 200 and 450 mM to prevent potential O_2_ diffusion limitation. Standardized instrumental and chemical calibrations were conducted according to the manufacturer’s instructions (Oroboros Instruments, Innsbruck, Austria). These allowed for corrections for i) background diffusion of O_2_ into the chamber, ii) O_2_ solubility in buffer Z, and iii) background consumption of O_2_ by the electrodes, which was determined across the range (200–450 mM) of chamber O_2_ concentrations employed during the experiments.

### Enzymatic activity

The maximal enzyme activity of citrate synthase (CS) was quantified in muscle homogenates, as previously described^107^. Enzymatic activity was normalized to g of total protein.

### Cell experiments

L6 myoblasts differentiation into myotubes was accomplished by reducing FBS concentration to 2% for 7 days and visually validated by phase contrast microscopy. 7-day differentiated L6-GLUT4*myc* myotubes seeded in 96-well plates were washed three times in pre-heated DPBS (Gibco, 14190144) and incubated for 4 h in BSA vehicle control solution (42 μM, i.e. the highest concentration added with palmitate), palmitate (250 μM), mitoquinone (50 nM), or combined palmitate and mitoquinone. DMSO (0.000001%) served as control for myotubes stimulated with mitoquinone. During the last 15 minutes of the incubation period, myotubes were stimulated with or without insulin (100 nM). Next, L6 myotubes were washed three times in ice-cold DPBS and fixed in 3% paraformaldehyde prior to blocking in 5% goat serum (Gibco, 16210064) and incubation with Myc-Tag antibody (1:500 in 5% goat serum). Subsequent incubation in HRP-conjugated secondary antibody allowed for colorimetric detection of plasma membrane-inserted, exofacially exposed GLUT4*myc* using the *o*-phenylenediamine method^126^.

C2C12 myoblasts differentiation into myotubes was initiated by switching to differentiation medium comprising DMEM supplemented with 2% horse serum (Cat. 26050-070; Thermo Fisher Scientific) and 1% penicillin-streptomycin. The differentiation medium was replaced every two days during the seven days of differentiation. 7-day differentiated myotubes maintained in 12-well-plates were stimulated with C2-ceramide (1-10 μM) or vehicle (0.1% DMSO) for 4 h, followed by incubation in NEM. C2C12 myotubes were homogenized in ice-cold lysis buffer (10% glycerol, 1% NP-40, 150 mM NaCl, 50 mM HEPES (pH 7.5), 20 mM sodium pyrophosphate, 10 mM NaF, 20 mM β-glycerophosphate, 2 mM phenylmethylsulfonyl fluoride, 1 mM EDTA (pH 8.0), 1 mM EGTA (pH 8.0), 2 mM Na3VO4, 10 μg/mL leupeptin, 10 μg/mL aprotinin, 3 mM benzamidine) containing 100 mM NEM. Each sample was sonicated for 15 s (10% intensity, Q2000 Qsonica sonicator) and the lysate supernatants were collected by centrifugation at 18,320 *g* for 10 min at 4°C.

### Animal experiments

#### In vivo gene transfer in mouse skeletal muscle

Skeletal muscle electroporation was performed in FDB muscles as previously described^65^. Male C57BL/6N mice 12-14 weeks of age were anesthetized by 2-3% isoflurane inhalation and injected into the foot sole with 10 μL of 0.36 mg/mL hyaluronidase (H3884, Sigma) diluted in PBS. After 60 min, 20 μL of sterile saline containing 20 μg of DNA was injected at the same location. Electroporation of the FDB muscle was performed by delivering 15 20-ms pulses at 1 Hz at an electrical field strength of 75 V/cm using acupuncture needles (0.20×25 mm, Tai Chi, Lhasa OMS) connected to an ECM 830 BTX electroporator (BTX Harvard Apparatus). After the procedure, the animals were transferred back to their cages and allowed to recover for at least 7 days.

#### Fiber isolation and GLUT4 translocation assay

Mice were sacrificed by cervical dislocation, and FDB was quickly removed under a dissection microscope. Muscles were incubated in serum-free alpha-Minimum Essential Medium Eagle (α-MEM) containing 2.0 mg/ml collagenase type 1 from Clostridium histolyticum (C0130, Sigma-Aldrich) for 2 h at 37°C. After collagenase treatment, muscles were incubated in 2 mL α-MEM containing 5% horse serum (26050-70, Gibco), followed by mechanical dissociation of individual muscle fibers using fire-polished Pasteur pipettes. Pooled fibers were seeded on 13 mm diameter glass coverslips (Hounissen) coated with Engelbreth-Holm-Swarm murine sarcoma ECM gel (E1270, Merck). After 1 h, 1 mL of α-MEM containing 5% fetal bovine serum was added to the wells, and the muscle fibers were cultured overnight. The following day, muscle fibers were maintained in serum-free α-MEM for 4 h containing either vehicle (0.1% ethanol) or 10 µM mitochondria-targeted paraquat (MitoPQ) and stimulated with either saline or 100 nM insulin for 20 min. Muscle fibers were fixed in 4% paraformaldehyde (4% PFA) diluted in PBS for 10 min. PFA-fixed muscle fibers were washed 3⊆10 min in PBS and incubated in blocking buffer [1% bovine serum albumin, 5% goat serum (16210-064; Gibco), and 0.1% sodium azide (247-852; Merck)] for 1 h. The muscle fibers were then incubated overnight with an antibody against the *myc* tag diluted 1:400 in blocking buffer. The next day, the fibers were washed 3×10 min in PBS containing 0.04% saponin and incubated with Alexa Flour 568 antibody in blocking buffer containing 0.04% saponin for 2 h. Finally, the fibers were washed three times, each for 10 min, in PBS and mounted on glass slides in Vectashield (H-1000; Vector Laboratories).

#### Live imaging procedures

Images were collected using a 40x 1.2 NA oil immersion objective on an LSM 980 confocal microscope (Zeiss) driven by Zeiss Zen Blue 3. To determine the fraction of GLUT4 accumulated on the plasma membrane surface, both GFP and *myc* signals were quantified as previously described^65^. Briefly, the *myc* and GFP channels were duplicated, and the new images underwent background subtraction and Gaussian blur adjustment (radius = 2) and were made binary. From these binary images, regions of positive GFP and *myc* signals were selected and overlaid onto the original images for quantification of the integrated density of GFP and *myc* signals, respectively.

### Quantification and statistical analysis

#### Calculations

Muscle glucose uptake and muscle lactate release, as measured by the net exchange of substrates (glucose and lactate) across the leg muscles, were calculated from the femoral arterial plasma flow (*F_p_*, expressed per kg of leg muscle mass) and the arteriovenous difference in plasma concentration of the given substrate (*substrate_a-v_*), which was corrected for changes in plasma volume by accounting for the net transcapillary water flux (*Jv*)^127, 128^:

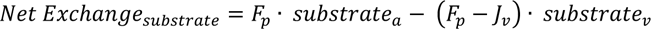

The femoral arterial plasma flow (*F_p_*) was calculated as:

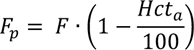

where *F* is the femoral arterial blood flow and *Hct_a_* is the arterial haematocrit.

The net transcapillary water flux (*J_v_*) into or out of the vein was calculated as:

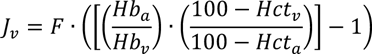

where *Hb_a_* and *Hb_v_* are the arterial and venous haemoglobin concentration, respectively, and *Hct_v_* is the venous haematocrit.

Leg O_2_ consumption 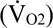, leg CO_2_ release 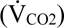 and leg respiratory quotient (RQ) were calculated from the femoral artery blood flow (F) and the arteriovenous difference in content of the given gas:

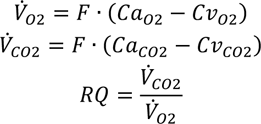

Content of O_2_ and CO_2_ in arterial blood (Ca_O2_ and Ca_CO2_) and venous blood (Cv_O2_ and Cv_CO2_) were computed using the equations by Siggaard-Andersen et al.^129^.

Glucose and lipid oxidation across the leg were estimated from the leg 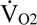 and 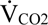 using the equations by Péronnet and Massicotte^130^.

### Statistics

For clamp-based outcomes, muscle insulin signaling and muscle redox state, between-treatment differences (“Lipid” vs. “Lipid + mtAO”) were determined using a linear mixed model including treatment as a fixed factor, subjects as a random factor, and with an unstructured covariance pattern. Within-treatment differences (i.e., Baseline vs. End-clamp or Pre-clamp vs. End-clamp) were determined using a linear mixed model with treatment-time interaction as a fixed factor, subject as a random factor, and with an unstructured covariance pattern. To adjust for the period effects associated with the crossover design of the study, a period variable was included as a fixed factor in the linear mixed models. For mitochondrial bioenergetics, between-treatment differences (“Control” vs. “P-CoA” vs. “P-CoA + mtAO”) were determined using a linear mixed model with treatment as a fixed factor and subjects as a random factor. For all analyses, model checking was based on Shapiro-Wilk’s test and quantile-quantile (Q-Q) plots. In case of heteroscedasticity (i.e., unequal variance), log transformation was applied before analysis. P values were evaluated using Kenward-Roger approximation of the degrees of freedom. Analyses were performed using SAS Enterprise Guide version 7.15.

For rodent muscle cell and fiber experiments, as well as for indeces of mitochondrial ADP sensitivity (EC_50_ and IC_50_) in human PmFBs, a one-way ANOVA in GraphPad Prism was used to estimate between-treatment differences.

The level of significance for all analyses was set at P < 0.05. Data are graphically presented as observed individual values with model-based estimated means ± 95% confidence limits, unless otherwise stated.

## SUPPLEMENTAL INFORMATION

**Table S1.**
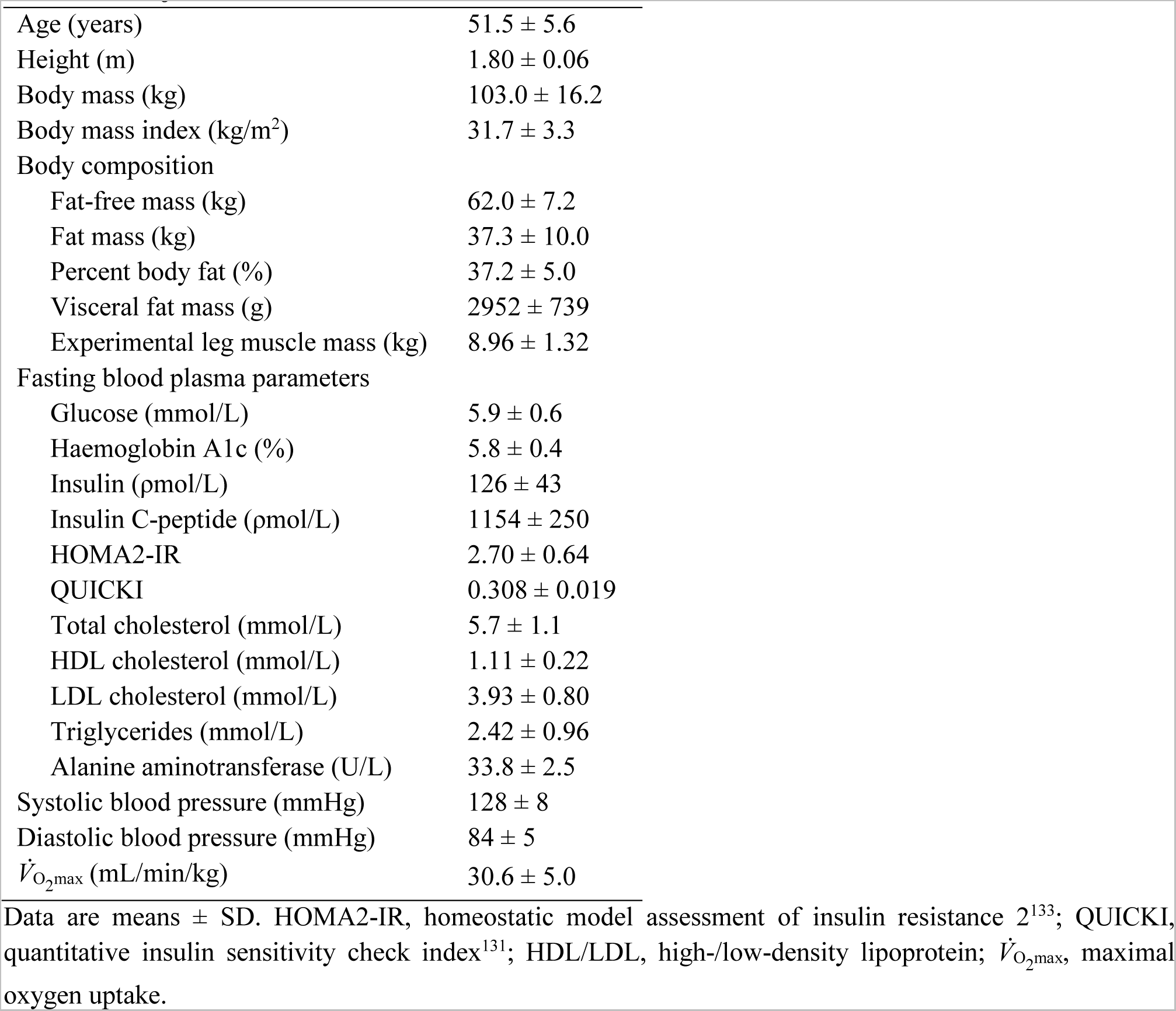
Subject characteristics.

**Figure S1.**
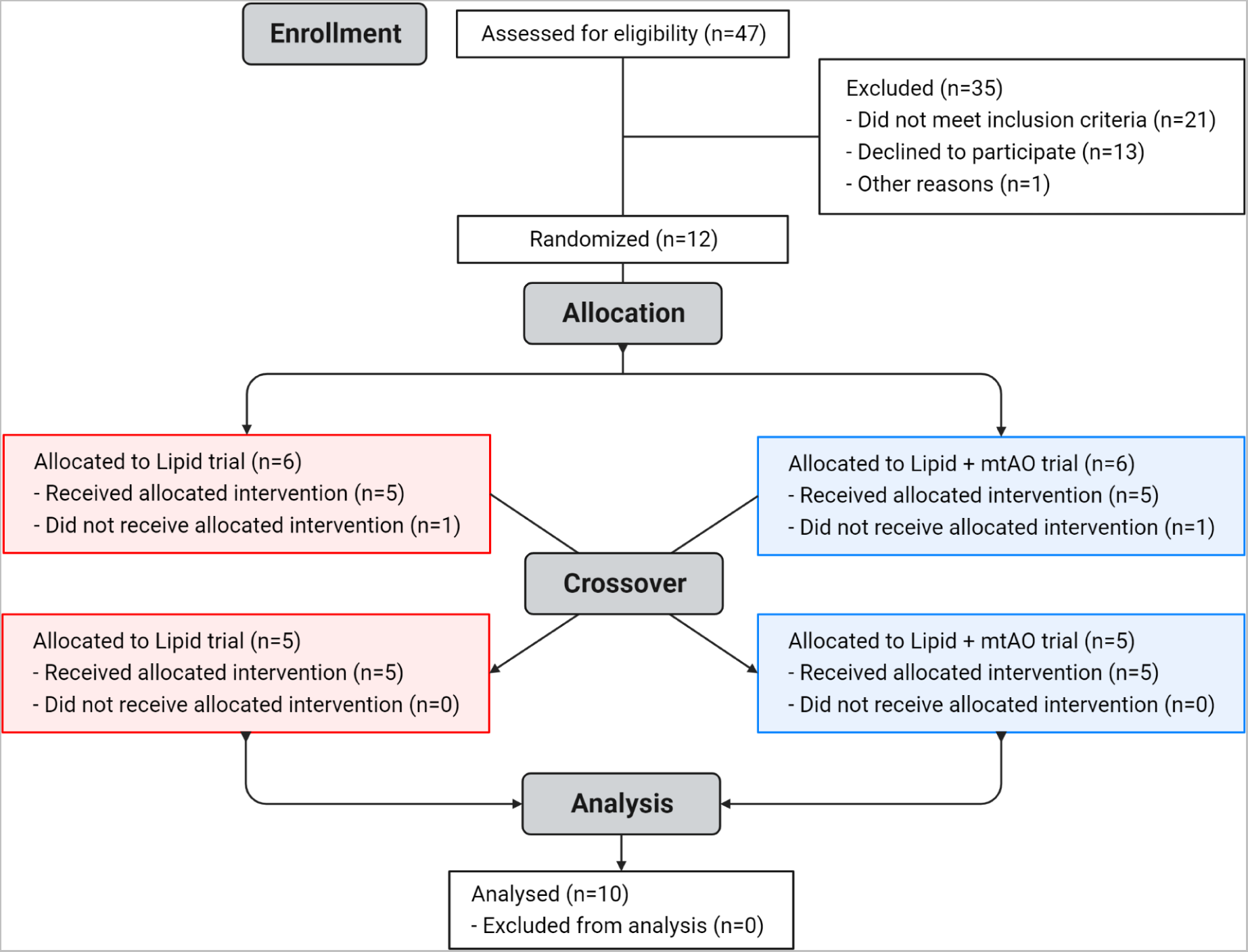
CONSORT flow diagram.

**Figure S2.**
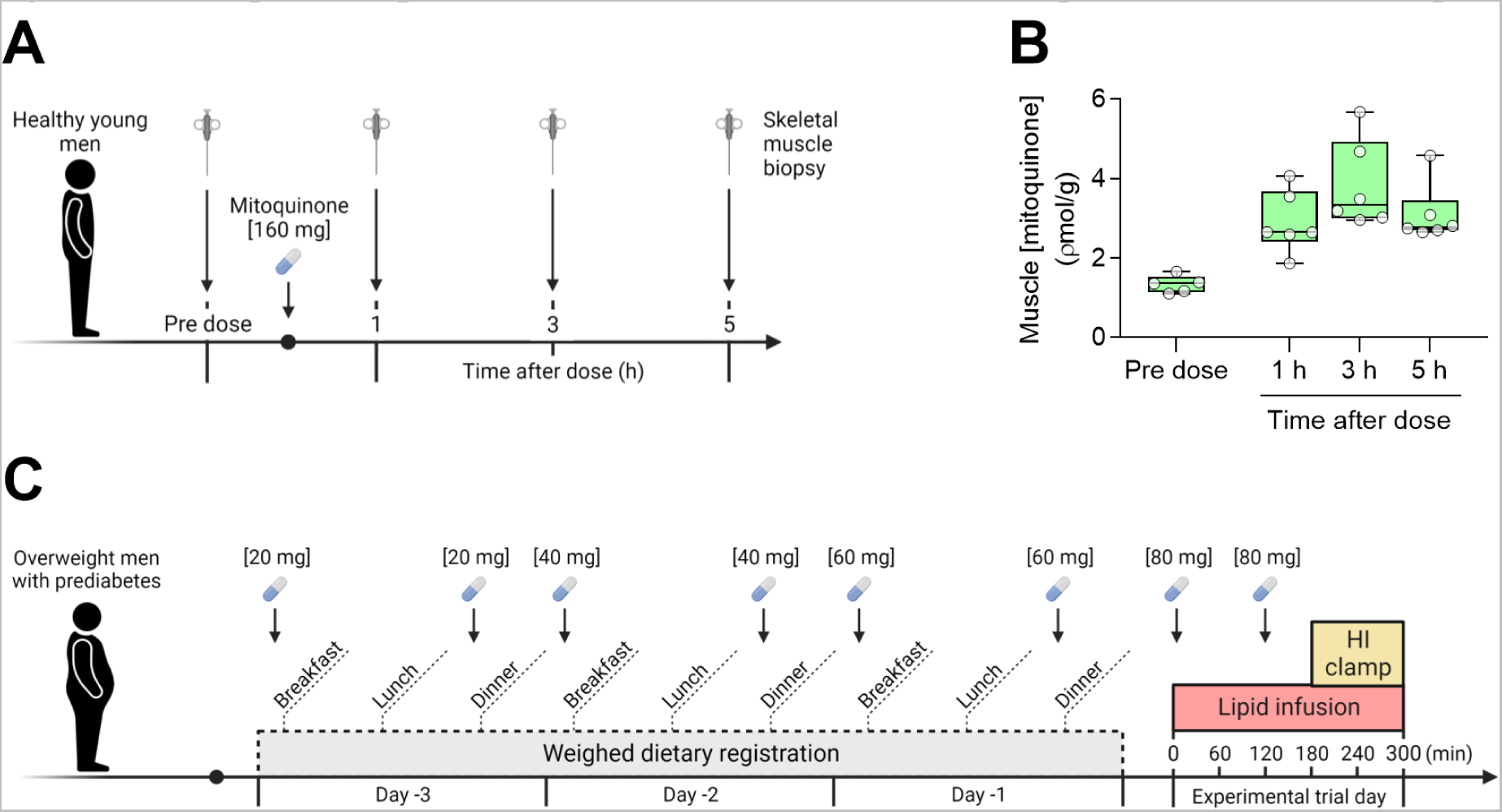
Mitoquinone uptake in human skeletal muscle and mitoquinone administration protocol. (A) Protocol employed in preliminary studies to evaluate pharmacokinetics of mitoquinone in skeletal muscle of healthy young men (n = 6). (B) Mitoquinone content in skeletal muscle, as measured by liquid chromatography-tandem mass spectrometry in muscle biopsy samples undertaken before and 1, 3 and 5 h after administration of a single oral dose [160 mg] in healthy young men. (C) Mitoquinone (or placebo) administration protocol employed in the main study.

